# Structural basis of substrate recognition and conformational gating in the bacteriophage M15 metalloendopeptidase LysPH

**DOI:** 10.64898/2026.04.21.719881

**Authors:** Dhruv Pahwa, Sakshi Pandey, Neelam Pandey, Kamble Amith Kumar, Priyanka Choudhary, Ardra Unni, Ishika Choudhary, Meet Ashar, Balram Ji Omar, Madhur Uniyal, Satyanarayan Rao, Ashwani Kumar Sharma, Shailly Tomar, Pravindra Kumar

## Abstract

The rapid emergence of antimicrobial resistance has led to a surge in multidrug-resistant strains, jeopardising the efficacy of frontline and last-resort antibiotics and thereby aggravating the antimicrobial resistance crisis. Bacteriophage-derived endolysins represent a promising class of next-generation antimicrobials. Here, we report the isolation of a novel bacteriophage, PA_Ganga_001, targeting multidrug-resistant *Pseudomonas aeruginosa*. Genomic sequencing of this phage identified a previously uncharacterised endolysin LysPH, a zinc-dependent globular endolysin that exhibits antibacterial activity against multiple multidrug-resistant Gram-negative pathogens, including *Pseudomonas aeruginosa, Acinetobacter baumannii*, and *Klebsiella pneumoniae*. Site-directed mutagenesis of the endolysin LysPH suggests that His77, Asp84, His159, Arg41, and Asp156 are essential for catalysis and substrate accommodation. To understand the structural basis of substrate binding and catalysis, we determined the crystal structures of the apo-enzyme (1.8 Å) and the catalytically attenuated D156A mutant in complex with a synthetic pentapeptide stem (PGX) of peptidoglycan (2.0 Å). The pentapeptide-bound complex revealed a defined substrate-binding groove, and a pronounced displacement of the Thr42-Ser75 loop was observed. This led to an approximately six-fold volumetric expansion of the catalytic cleft, indicative of a substrate-induced transition to an open, catalytically competent conformation. Notably, a distinctive C-terminal helical element, which diverges sequentially from characterised M15 homologues, is predicted to interact with the NAG-NAM scaffold and may contribute to the positioning of the stem pentapeptide for Zn^2+^ dependent catalysis. These distinct structural features establish the molecular basis for L-Ala-D-Glu hydrolysis by M15 bacteriophage endolysins and provide a foundational framework for the rational design and engineering of the next generation of antimicrobial enzymes.

## Introduction

The rise of multidrug-resistant bacteria is a major threat to global health, severely limiting the effectiveness of current antibiotic treatments. The ESKAPE pathogens include *Pseudomonas aeruginosa, Acinetobacter baumannii, Klebsiella pneumoniae, Enterococcus faecium, Staphylococcus aureus*, and *Enterobacter spp*. These bacteria are associated with considerable morbidity and mortality and have high levels of antimicrobial resistance, underscoring the need to develop alternative antimicrobial strategies. Among ESKAPE pathogens, *P. aeruginosa* is particularly problematic, causing a range of nosocomial, acute, and chronic infections and resulting in over 500,000 deaths worldwide annually, with more than 300,000 cases attributed to antimicrobial resistance.^1–3^

Bacteriophage-derived endolysins are peptidoglycan hydrolases which cleave specific bonds within the bacterial peptidoglycan and are classified as glycosidases, amidases, and endopeptidases, each targeting specific glycosidic or peptide linkages within the peptidoglycan network.^4,5^ Owing to their rapid bactericidal activity and low propensity to develop resistance, endolysins are being explored as alternative therapeutics against multidrug-resistant pathogens, including topical wound treatment and engineered variants with enhanced stability and broader host specificity. Among these, L-alanyl-D-glutamate peptidases of the M15 family of metalloendopeptidases are zinc-dependent enzymes that hydrolyse the L-Ala–D-Glu bond within the peptidoglycan stem peptide through Zn^2+^-activated catalysis.^6–8^

Despite advances in the structural characterisation of endolysins, mechanistic insight into M15 metalloendopeptidases remains limited. While peptidoglycan-bound structures have been reported for modular muramidase endolysins, such as Cpl-1, comparable structural information is lacking for globular M15 endopeptidases. Notably, no crystal structure is available for a bacteriophage-derived globular M15 endopeptidase bound to an intact stem pentapeptide, and the molecular basis of how the glycan backbone and L-Ala–D-Glu bond are positioned within the catalytic Zn^2+^ site remains poorly defined. Furthermore, although some Gram-negative endolysins harbour C-terminal helices, the exact role of these helices in substrate recognition and positioning remains unclear. ^9–11^

Recently, the crystal structures of bacteriophage-derived M15 endolysins have provided insights into the conserved elements and functional diversity of M15 endolysins. The crystal structure of LysB4 (PDB: 6AKV) established a canonical LAS-type Zn^2+^-dependent active-site architecture, providing a framework for L-alanyl–D-glutamate endopeptidase activity. Similarly, structures of the T5 endolysin EndoT5 (PDB: 2MXZ, 8P3A) have revealed extended loop regions that undergo conformational stabilisation upon binding of a regulatory Ca^2+^ ion, suggesting a dynamic gating mechanism that modulates substrate access to the catalytic centre. In addition, LysECD7 (PDB: 8Q2G), which shares substantial sequence similarity with LysPH, has been engineered by incorporating C-terminal amphipathic helices to enhance activity against Gram-negative bacteria. Despite these advances, structural insight into how peptidoglycan substrates are positioned within the catalytic cleft of globular M15 endolysins remains limited, and the contribution of non-conserved C-terminal elements to substrate recognition has not been clearly defined.^12–14^

Here, we characterise LysPH, a zinc-dependent M15 endopeptidase encoded by the bacteriophage PA_Ganga_001. LysPH specifically hydrolyses the L-Ala–D-Glu bond within the peptidoglycan stem peptide via Zn^2+^-activated catalysis and exhibits lytic activity against multiple Gram-negative pathogens, including *P. aeruginosa, A. baumannii* and *K. pneumoniae*. Crystal structures of the native enzyme and the D156A mutant in complex with a stem pentapeptide substrate, together with structure-guided mutagenesis, define the molecular basis of substrate recognition and catalysis. By successfully capturing the enzyme in both a substrate-free ‘closed’ state and a pentapeptide-bound ‘open’ conformation, we reveal a well-defined binding groove and a pronounced displacement of the Thr42-Ser75 loop, consistent with a substrate-induced conformational gating mechanism. We identify a C-terminal helix, which diverges sequentially from characterised M15 homologues, and is predicted to position the peptidoglycan and the stem pentapeptide within the catalytic Zn^2+^ site. We propose that the Thr42-Ser75 loop and this C-terminal structural element act cooperatively to mediate substrate alignment and access to the catalytic centre, thereby regulating enzyme activity.

## Results

### Phage PA_Ganga_001 encodes the previously uncharacterised endolysin LysPH

The endolysin LysPH was identified within the lysis cassette of the novel bacteriophage PA_Ganga_001. This phage was isolated from an environmental sample collected from the Ganga River in Rishikesh (30°06′30″N, 78°17′50″E), and its genome was sequenced and deposited in GenBank (accession no. PP597238). PA_Ganga_001 was found to encode three proteins involved in the lysis cassette: a putative holin, an Rz-like spanin, and a putative endolysin LysPH. Using BLASTp analysis against the NCBI non-redundant database, LysPH was classified within the MEROPS M15 family of metallopeptidases. These proteins are zinc-dependent metallopeptidases that hydrolyse peptide cross-links within bacterial peptidoglycan, suggesting a role in peptidoglycan cleavage during host lysis. (Fig. S1)

### Structural and functional characterisation of LysPH

Recombinant LysPH was cloned, expressed in *E. coli*, and purified to homogeneity using affinity and size-exclusion chromatography. The purified enzyme exhibited a monomeric behaviour in solution, consistent with other globular M15 endopeptidases.

The crystal structure of LysPH was determined at 1.8 Å resolution and belongs to the P2_1_ 2_1_ 2_1_ space group, with a single monomer in the asymmetric unit. The structure was solved by molecular replacement using the LysECD7 structure (PDB: 8Q2G) as the search model. LysPH has a single globular domain with an (α/β)_4_ topology. A prominent loop–helix segment spanning residues Thr42–Ser75 forms a flexible flap-like element that partially covers the active site, showing a closed conformation.

Structural comparison with other MEROPS M15 family members identified a zinc-dependent endopeptidase fold. The catalytic site is defined by three residues that coordinate a Zn^2+^ ion (His77, Asp84, and His159), with Asp156 positioned close to the Zn^2+^-bound water molecule. Residues Arg41 and Ser55 line the active-site cleft and contribute to substrate binding, forming a charged groove compatible with peptidoglycan recognition. (Fig. 1A, B) PDB-BLAST search revealed that LysPH shares 42% sequence identity with endolysin LysECD7 (PDB: 8Q2G) and 36% with bacterial peptidoglycan hydrolase ChiX (PDB: 5OPZ), a characterised L-alanyl-D-glutamate endopeptidase. Consistently, DALI searches showed strong structural homology with bacteriophage endolysins LysB4 from Bacillus cereus phage B4 (PDB: 6AKV) and EndoT5 from Escherichia phage T5 (PDB: 2MXZ), both of which are known L-alanyl-D-glutamate endopeptidases. The catalytic active site was conserved among these homologs, suggesting that LysPH functions as a zinc-containing L-alanyl-D-glutamate endopeptidase that cleaves the bond between L-alanine and D-glutamate in the peptidoglycan stem pentapeptide. (Fig. 1C, D)^12–14^

**Figure 1.**
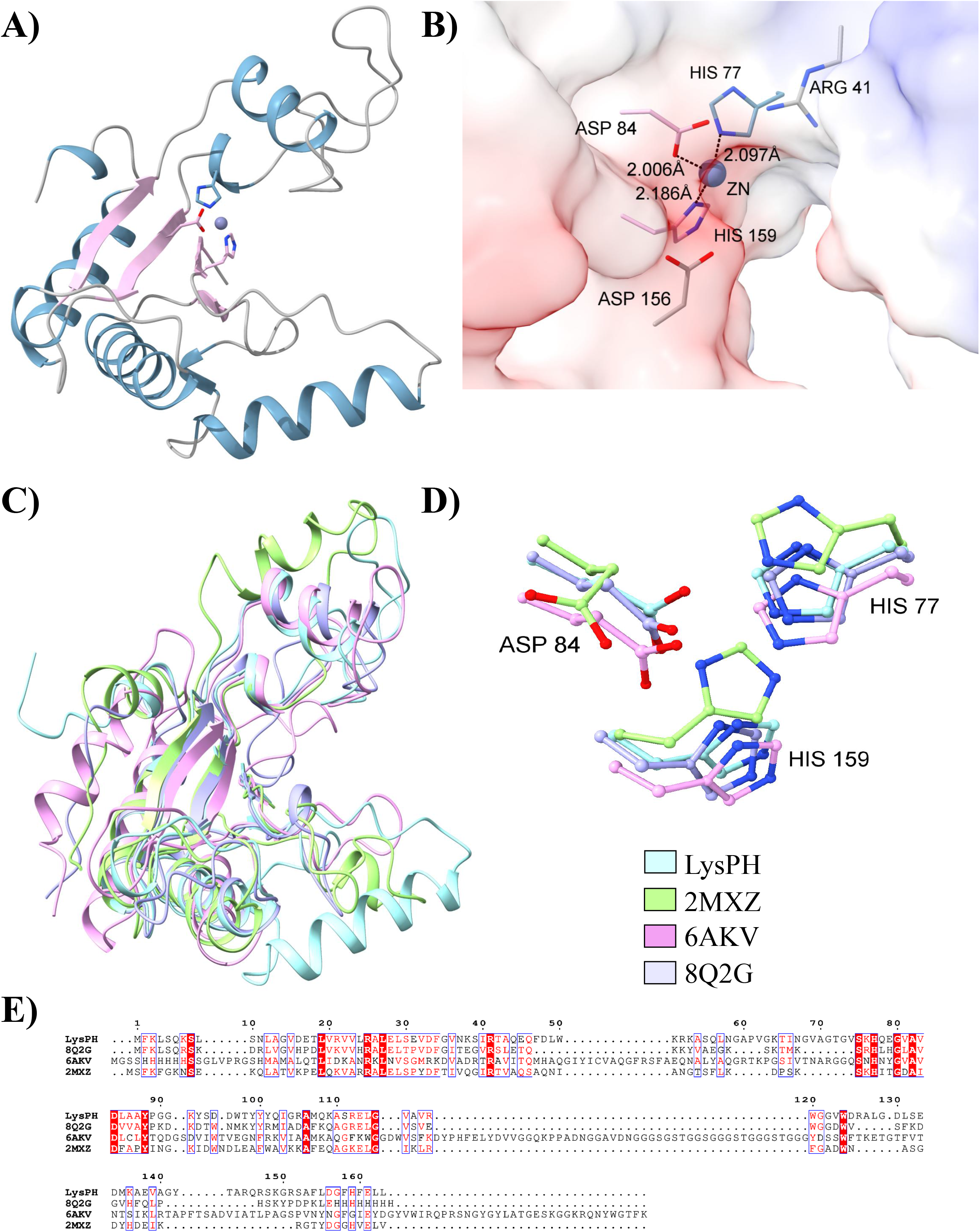
High-resolution crystal structure of apo-LysPH establishes the conserved M15 metalloendopeptidase fold. (A) Crystal structure of LysPH endolysin (PDB: 21UA). (B) Electrostatic surface representation of the substrate-binding groove. The Zn^2+^ ion (grey sphere) is coordinated by His77, His159, and Asp84, with Arg41 and Asp156 positioned near the catalytic centre. Selected metal–ligand and inter-residue distances are indicated in Å. The electrostatic surface is coloured from ™5 (red) to +5 (blue) kBT/e. (C) Structural superposition of LysPH with its closest structural homologues (PDB: 8Q2G, 6AKV, and 2MXZ), showing conservation of the overall fold. (D) Close-up comparison of the catalytic residues of LysPH with those of homologous structures, highlighting the conserved arrangement within the active site. (E) Multiple sequence alignment of LysPH with representative M15 endopeptidase homologues.

### Antibacterial activity of LysPH and its mutants

The functional roles of residues involved in catalysis and substrate binding were investigated using alanine-substitution mutants (R41A, S55A, H77A, D84A, H159A, and D156A), which were analysed by size-exclusion chromatography (SEC) to assess their structural integrity. Wild-type LysPH eluted as a single, symmetric peak (V_e_ = 106.62 mL), consistent with a monodisperse monomer. Five mutants (R41A, H77A, D84A, H159A, and D156A) exhibited SEC profiles comparable to wild-type, with minimal shifts in elution volume (ΔV_e_ ≤ 1.54 mL), similar peak widths, and no evidence of aggregation, indicating preservation of the global fold and oligomeric state. The S55A mutant displayed a markedly aberrant profile (V_e_ = 120.84 mL, K_av_ = 1.010), with a broadened, asymmetric peak, indicative of structural perturbation and conformational heterogeneity. Together, these results demonstrate that most active-site mutations do not affect protein folding, whereas substitution of Ser55 compromises structural integrity.

A comprehensive dose-response analysis was performed to quantify the specific potency of wild-type LysPH and establish an optimal concentration for comparative mutant screening. The enzyme exhibited concentration-dependent bacteriolytic activity with a sharp transition phase. Fitting the endpoint lysis data at 30 minutes to a four-parameter logistic (4PL) model yielded an EC50 of 6.00 µM and a Hill coefficient of 3.01. Notably, the lytic efficacy jumped from 6.81 ± 2.88% at 3 µM to a near-maximal 60.89 ± 4.99% at 10 µM. Because 10 µM is the lowest concentration tested that reliably yields robust, near-maximal lysis, it was selected as the standard baseline for all subsequent mutational profiling. (Fig. 2A, B)

**Figure 2.**
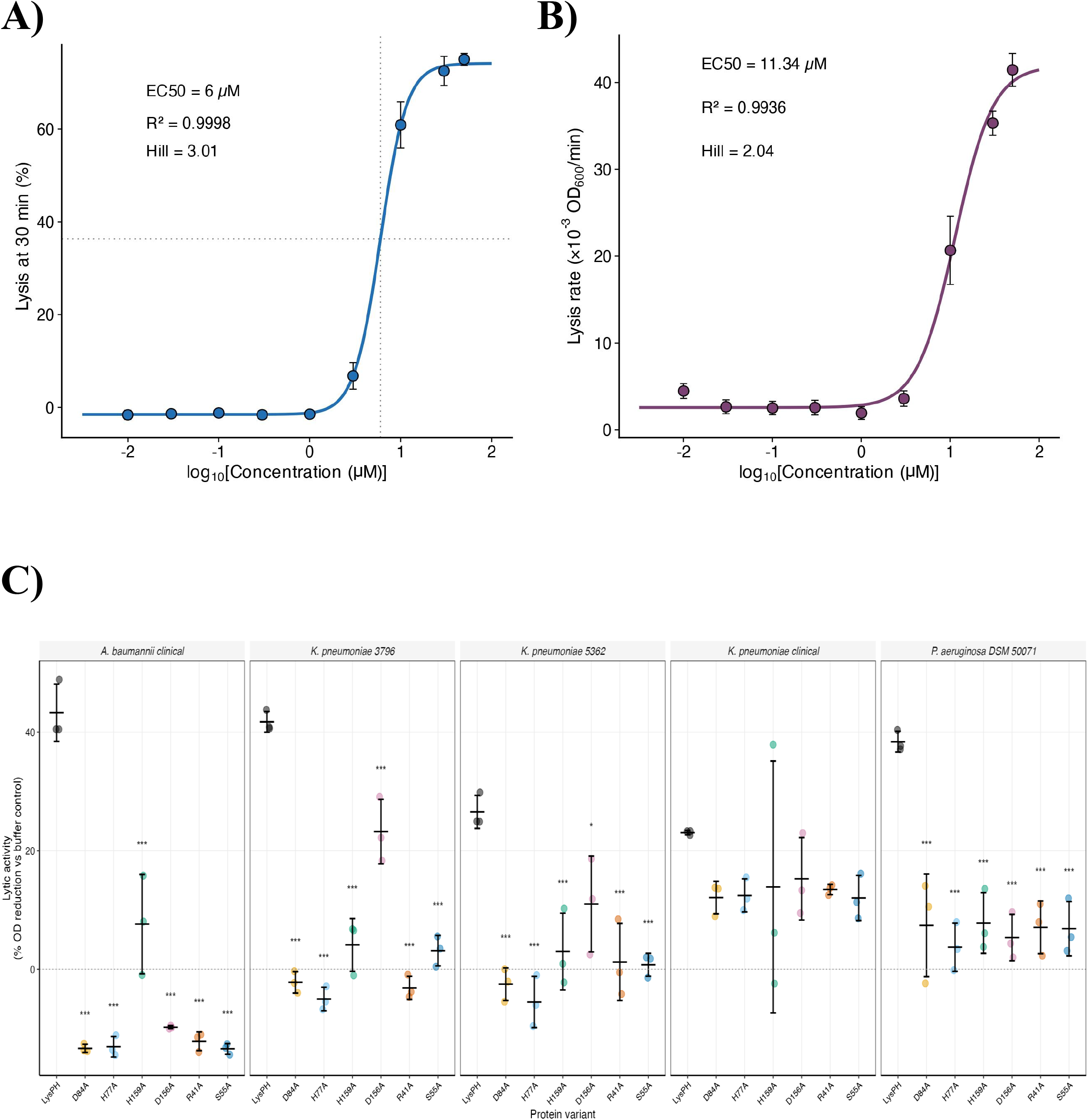
Dose–concentration profiling of wild-type LysPH and comparative lytic activity of active-site variants. (A) Concentration-dependent lytic activity of wild-type LysPH, plotted as endpoint % *OD*_*600*_ reduction after 30 min. Data were fitted to a four-parameter logistic (4PL) model, yielding an EC_50_ of 6.0 µM and a Hill coefficient of 3.01. (B) Concentration-dependent initial lysis rates of wild-type LysPH, calculated by linear regression over the first 20 min, with 4PL fitting yielding a rate-based EC_50_ of 11.3 µM and a Hill coefficient of 2.04. Data in panels A and B represent mean ± SD from three independent biological replicates. (C). Lytic activity of LysPH and mutant variants at 10 µM, quantified as % OD_600_ reduction relative to buffer control over 60 min (AUC-based). Individual biological replicates are shown as colored dots (n = 3); horizontal bars and whiskers represent mean ± SD. Significance versus wild-type LysPH was assessed by Welch’s t-test with Holm correction for multiple comparisons (* p < 0.05, ** p < 0.01, *** p < 0.001). Wild-type LysPH showed robust lytic activity (23–43% OD reduction) across all five strains. Catalytic mutants D84A and H77A showed the most complete loss of function, while the non-catalytic mutant D156A retained partial activity in a strain-dependent manner.

The lytic spectrum of LysPH was characterised by monitoring OD_600_ reduction over 60 minutes against five Gram-negative and two Gram-positive bacterial strains at a uniform protein concentration of 10 µM. This concentration was selected based on measurements across a range of LysPH concentrations, which revealed a sharp transition in activity, with minimal lysis at 3 µM (6.8%) and a marked increase at 10 µM (∼60.9%). As this lies just above the EC_50_ (∼6.0 µM), it ensures robust wild-type activity while preserving sufficient dynamic range to sensitively detect residual activity in mutant variants. Wild-type LysPH produced rapid and substantial OD reduction in all Gram-negative strains tested, achieving 43.3 ± 4.8% reduction against *A. baumannii* clinical, 41.7 ± 1.8% against *K. pneumoniae 3796*, 26.6 ± 2.8% against *K. pneumoniae 5362*, 23.1 ± 0.4% against *K. pneumoniae clinical*, and 38.4 ± 1.7% against P. *aeruginosa DSM 50071*. The most pronounced lytic kinetics were observed against *A. baumannii*, where >50% of the total OD reduction occurred within the first 20 minutes (Fig. S3B, Table S1).

The roles of residues involved in catalysis and substrate positioning within the active site were assessed by measuring the lytic activity of all mutants using turbidity reduction assays. All five mutants showed markedly attenuated lytic kinetics compared to wild-type, with curves largely overlapping the buffer control in most strains. To quantify the magnitude of lytic impairment, we calculated % OD reduction relative to control using area-under-curve (AUC) analysis (Fig. 2C, Table S2).

Alanine substitution of the three Zn^2+^-coordinating residues resulted in severe loss of activity across all strains. D84A showed near-complete abolition of lytic function, with mean % OD reduction of −13.3 ± 0.7% in *A. baumannii* (relative activity −0.31 vs WT) and −2.2 ± 1.8% in *K. pneumoniae 3796* (relative activity −0.05; p < 0.001). H77A exhibited a comparable profile, with relative activity of −0.30 in *A. baumannii* and −0.12 in *K. pneumoniae 3796* (p < 0.001). H159A, the third catalytic residue, showed severely reduced but detectable residual activity (relative activity 0.10–0.20 across strains; p < 0.01).

The D156A mutant exhibited a different activity profile compared to the catalytic mutants. D156A retained substantial lytic activity against *K. pneumoniae 3796* (23.2 ± 5.4% OD reduction; relative activity 0.56; p < 0.05) and *K. pneumoniae 5362* (11.0 ± 8.1%; relative activity 0.41; p = 0.067, ns after Holm correction). In contrast, D156A showed near-complete loss of activity against *A. baumannii* (−9.8 ± 0.3%; relative activity −0.23; p < 0.01) and markedly reduced activity against *P. aeruginosa* (5.3 ± 3.9%; relative activity 0.14; p < 0.01).

Based on structural comparison with homologous M15 endopeptidases and the positioning of Asp156 in proximity to the catalytic Zn^2+^-bound water molecule, it was initially hypothesised to function as a general base involved in water activation during catalysis. However, the retention of residual activity in the D156A mutant indicates that Asp156 is not essential for catalysis and instead likely plays a role in substrate positioning and optimal alignment of the scissile bond within the active site.

Alanine substitution of the substrate-binding residue Arg41 resulted in a marked loss of lytic activity comparable to that observed for catalytic mutants. R41A showed relative activity of −0.28 in *A. baumannii* (p < 0.01), −0.07 in *K. pneumoniae 3796* (p < 0.001), and 0.05 in *K. pneumoniae 5362* (p < 0.05), demonstrating that disruption of substrate binding severely impairs enzymatic function.

The S55A mutant exhibited a similarly reduced activity profile (relative activity −0.31 in *A. baumannii* (p < 0.01), 0.08 in *K. pneumoniae* 3796 (p < 0.001), and 0.03 in *K. pneumoniae* 5362 (p < 0.01)); however, interpretation of this effect is complicated by evidence of structural perturbation observed in size-exclusion chromatography.

The addition of protein to the assay well may contribute to baseline absorbance through light scattering or refractive index effects, resulting in apparent OD values marginally above the buffer control, as the magnitude (<15%) was consistent across inactive mutants.

### Crystallisation of LysPH_D156A complexed with peptidoglycan pentapeptide

The D156A mutant was selected to capture the enzyme–substrate complex, based on its proximity to, but not direct involvement in, Zn^2+^ coordination. Substitution of Asp156 was therefore expected to reduce catalytic turnover while preserving the integrity of the metal-binding site, thereby facilitating substrate trapping. The crystal structure of LysPH_D156A was determined at a resolution of 2Å. The crystals belonged to the P2_1_ 2_1_ 2_1_ space group, with a single monomer in the asymmetric unit, and the structure was obtained by molecular replacement using the LysPH structure (PDB: 21UA) as the search model.

The geometry of the catalytic Zn^2+^ coordination sphere was preserved in both the apo and substrate-bound structures. In the wild-type LysPH structure, the Zn^2+^ ion is coordinated by His159, His77, and Asp84 at distances of 2.2 Å, 2.1 Å, and 2.0 Å, respectively (Fig. 3). In the LysPH_D156A–pentapeptide complex, these distances remain largely unchanged (2.1 Å, 2.0 Å, and 2.0 Å, respectively), indicating that substitution of Asp156 does not perturb the primary metal coordination geometry. The L-alanine of the bound pentapeptide is in close proximity to the Zn^2+^ ion (∼2.1 Å), indicating that the substrate is in the correct orientation in the catalytic site.

**Figure 3.**
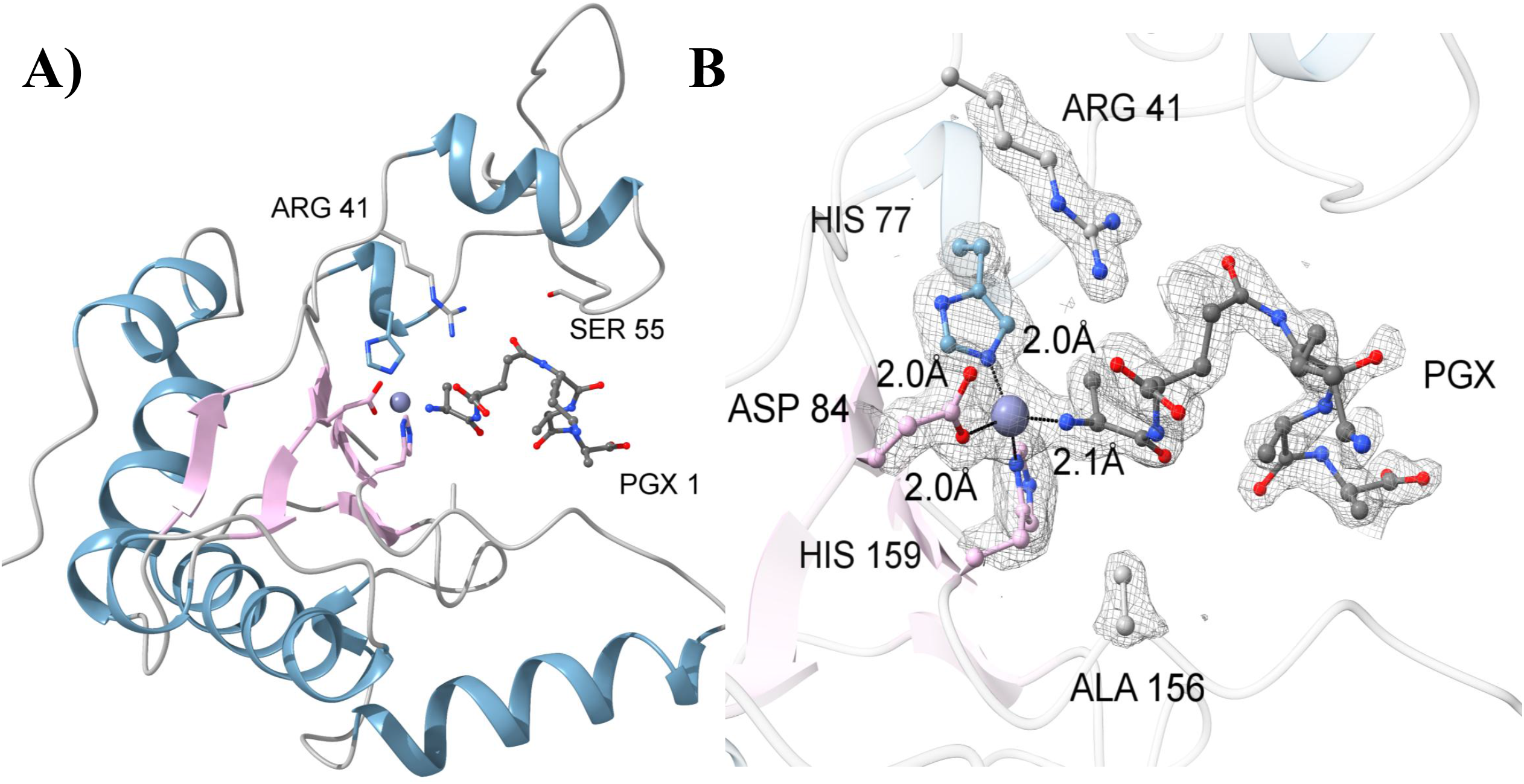
Capture of the LysPH-substrate complex reveals a substrate-induced gating mechanism. (A) Crystal structure of the D156A mutant of LysPH (PDB: 23VM) in complex with a PGX pentapeptide (L-Ala–D-iGlu–L-Lys–D-Ala–D-Ala) shown as orange sticks. Structure of the wild-type enzyme used for comparison (PDB: 21UA). (B) Polder map (mFo–DFc, contoured at 3.0 σ; grey mesh) confirming electron density for the PGX ligand. Catalytic residues His^77^, Asp^84^, and His^159^, and substrate residues are labelled. (C) Superposition of selected frames from morph interpolation (frames 1, 10, 20, 30, 40, and 50) illustrating progressive displacement of the Thr42–Ser75 loop. Intermediate conformations are shown as semi-transparent representations to depict the trajectory between LysPH and LysPH_D156A, consistent with a conformational gating mechanism.

### Thr42–Ser75 flap dynamics facilitate substrate access

Comparing the apo LysPH and LysPH_D156A-pentapeptide complex structures, it is evident that although the overall backbone conformation is similar (Cα RMSD ∼0.56 Å), the Thr42-Ser75 loop, which constitutes part of the wall of the catalytic cleft, displays significant positional displacement upon binding of the substrate. The loop exhibits significant rigid-body motion, with an internal RMSD of 0.29 Å, and also exhibits significant movement of its centroid by ∼5.9 Å and the tip of the loop, Ser68, by ∼9.4 Å, thus expanding the catalytic cleft to accommodate the substrate. DynDom6D analysis identified two dynamic domains, with the Thr42–Ser75 loop forming part of a mobile domain (273 atoms) that undergoes rigid-body motion relative to a largely fixed core domain (937 atoms). This motion is characterised by a rotation of 32.5° accompanied by negligible translation (∼0.5 Å), indicating a hinge-like movement. The high interdomain-to-intradomain displacement ratio (2.08) confirms that this conformational change is dominated by rigid-body rotation rather than local flexibility, supporting a gating mechanism for substrate access.^15^ Furthermore, pocket volume calculations using CASTp confirmed that this rigid-body hinge motion triggers a massive expansion of the active-site cavity. The solvent-accessible volume of the catalytic cleft increases nearly six-fold, expanding from 108.8 Å^3^ in the closed apo-enzyme to 652.5 Å^3^ in the open substrate-bound state, thereby generating the necessary spatial dimensions to accommodate the bulky peptidoglycan stem pentapeptide.^16^ These observations indicate that the Thr42–Ser75 loop acts as a dynamic gating element that undergoes conformational rearrangement, regulating access to the catalytic site. (Fig. 4)

**Figure 4.**
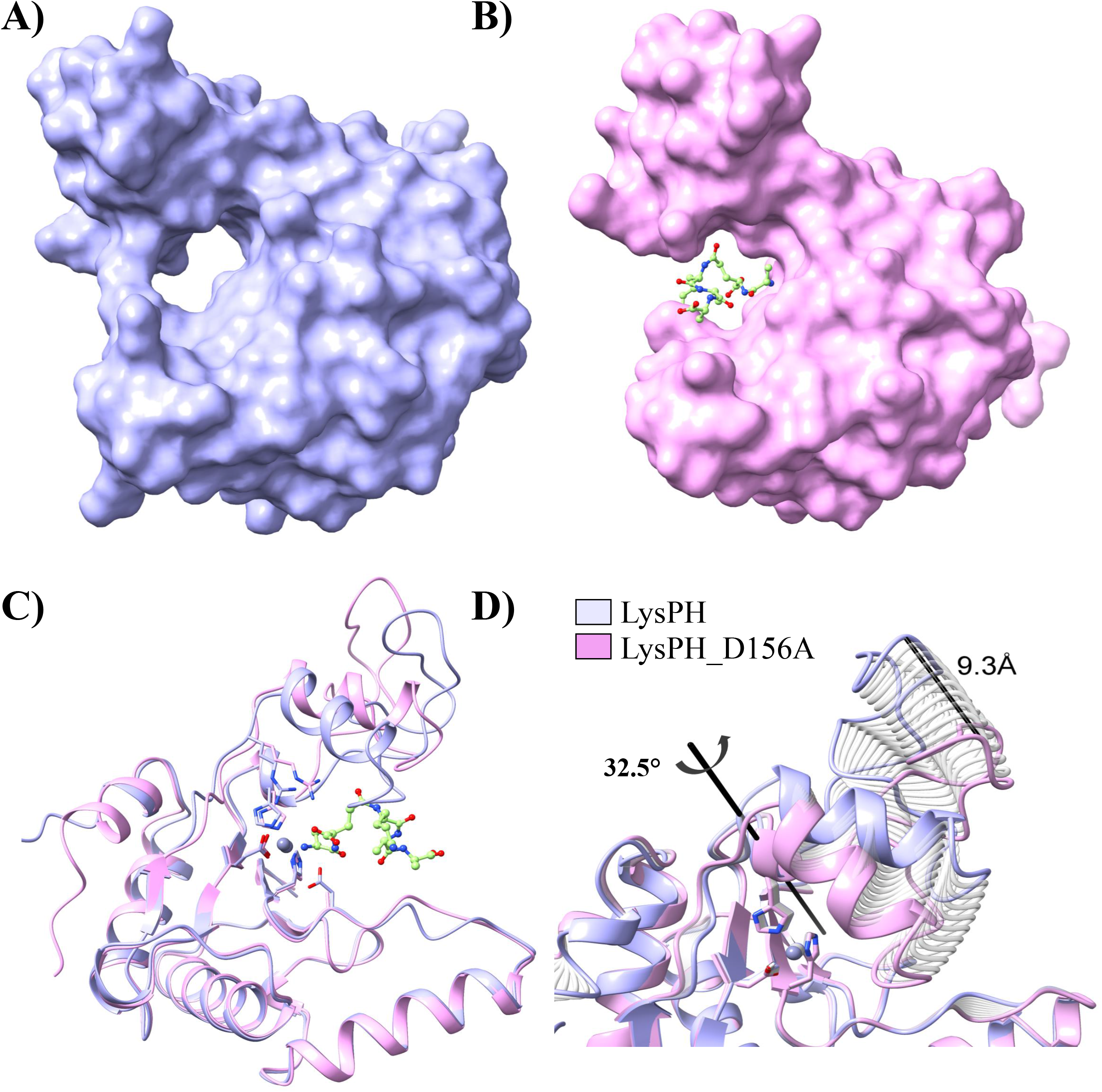
Substrate-induced conformational gating in LysPH endolysin(A) Surface representation of wild-type LysPH showing the closed conformation of the active-site cleft, with the Thr42–Ser75 loop occluding substrate access. (B) Surface representation of the LysPH_D156A–pentapeptide complex illustrating the open conformation, in which displacement of the Thr42–Ser75 loop exposes the catalytic groove to accommodate the substrate. (C) Structural superposition of LysPHD156A–PGX complex (Pink) with LysPHWT (Purple), highlighting movement of the substrate-binding flap. (D) Active-site structural rearrangement highlighting the Thr42–Ser75 loop trajectory. Semi-transparent morph frames reveal a 9.3 Å displacement at the Ser68 tip. DynDom6D analysis confirms a 32.5° rigid-body rotation of the mobile domain around the interdomain screw axis (cylinder) to expand the catalytic cleft.

### C-terminal helix as a phage-specific scaffold for substrate positioning

Structural superposition of LysPH with representative M15 family homologues (PDB: 2MXZ, 8Q2G, 6AKV) revealed that the C-terminal helix in LysPH is not conserved with structurally characterised homologues analysed here (Fig. 1C). Multiple sequence alignment showed a lack of strict conservation in this region (Fig. 1E), supporting the notion that the C-terminal extension represents a phage-specific structural adaptation.

Helical wheel analysis showed that the C-terminal helix lacks classical amphipathic character, as reflected by its low hydrophobic moment, high polarity, and absence of a defined hydrophobic face. These features suggest that helix does not play a primary role in membrane insertion, but instead supports a more specialised structural role in substrate positioning and structural support within the peptidoglycan-binding groove.^17^

To further investigate the role of the extended C-terminal region in a more physiologically relevant context, further MD simulations were performed using the minimal glycan–pentapeptide complex to represent the peptidoglycan substrate (NAM–NAG–NAM–L-Ala–D-Glu–L-Lys). The simulations suggested that although the C-terminal helical segment itself does not directly contact the NAG–NAM disaccharide, residues within the adjacent C-terminal loop extend toward the catalytic cleft, where they may interact with the glycan backbone and thereby orient the attached stem pentapeptide such that the scissile L-Ala–D-Glu bond is positioned proximal to the Zn^2+^ catalytic centre, facilitating alignment of the scissile bond. At the same time, Asp156 remains appropriately oriented to support catalysis. Notably, substitution of Phe154 resulted in a marked increase in local flexibility of the C-terminal region (residues 133–163), with the mean RMSF increasing from 0.39 Å in the wild-type complex to 0.79 Å in the F154A mutant, indicating localised destabilisation of the C-terminal segment (Fig. S5A, B).

Based on these observations, we propose a model in which the C-terminal helix serves as a scaffold, positioning the adjacent loop in an optimal conformation for substrate binding, thereby likely facilitating alignment of the L-Ala–D-Glu bond in the stem pentapeptide within the catalytic site.

### Data Deposition

The structures have been deposited in the Protein Data Bank with accession codes 21UA and 23VM and the genome of phage PA_Ganga_001 are submitted under GenBank accession no. PP597238

## Discussion

Bacteriophage-derived endolysins have been identified as promising candidates for the development of new antibacterials, especially against MDR pathogens, due to their capacity to hydrolyse highly conserved bonds in bacterial peptidoglycan. In this study, we identify and characterise PA_Ganga_001, a lytic Caudoviricetes phage infecting multidrug-resistant *P. aeruginosa*, and provide structural and mechanistic insights into its endolysin, LysPH. PA_Ganga_001 encodes a canonical holin–spanin–endolysin lysis cassette, and its endolysin LysPH is a zinc-dependent M15 metallopeptidase predicted to function as an L-alanyl–D-glutamate endopeptidase with activity against several multidrug-resistant Gram-negative pathogens. ^18,19^

### LysPH as an L-alanyl–D-glutamate M15 endopeptidase

Sequence and structural analyses indicate that LysPH belongs to the MEROPS M15 family of metallopeptidases. The enzyme adopts the conserved α/β fold characteristic of this family and contains a catalytic Zn^2+^ ion coordinated by His77, Asp84, and His159. Structural similarity to characterised endopeptidases such as LysECD7, LysB4, and EndoT5, together with conservation of active-site architecture, supports the conclusion that LysPH most likely functions as an L-alanyl–D-glutamate endopeptidase that cleaves the stem peptide of peptidoglycan.^12–14^

Nearly complete loss of activity upon alanine substitution of the Zn^2+^-coordinating His77 and Asp84 supports their role as essential residues required for metal coordination and catalysis. In contrast, the H159A mutant retained partial activity, suggesting that His159 contributes to maintaining optimal Zn^2+^ coordination geometry, which can still be partially maintained by His77 and Asp84.

In addition to catalytic residues, substitution of the substrate-binding residue Arg41, which lines the active-site cleft, resulted in a loss of activity comparable to that observed for catalytic-site mutants, indicating a critical role in substrate positioning within the active site. Substitution of Ser55 also caused an almost complete loss of enzyme activity. Overall, these findings suggest that precise alignment of the peptidoglycan pentapeptide within the active-site groove is crucial for attaining a catalytically active enzyme conformation, consistent with the established mechanism of peptidoglycan hydrolases.

### Thr42–Ser75 flap mediates substrate access via an induced-fit mechanism

Structural comparison of the apo enzyme and the pentapeptide-bound LysPH_D156A complex indicates that the Thr42–Ser75 loop–helix element, which forms one wall of the catalytic cleft, undergoes a pronounced hinge-like displacement upon binding of the substrate. In the apo structure, this loop adopts a closed conformation, partially occluding the active site, whereas in the substrate-bound complex, it transitions to an open conformation that exposes the catalytic cleft. Kinematic analysis confirms that the preservation of the loop’s internal conformation during this transition is a true rigid-body rotation, rather than localised flexibility or loop melting.

This substrate-induced hinge motion results in a massive structural expansion of the active-site cavity. Pocket volume calculations show a nearly six-fold increase in the solvent-accessible volume of the catalytic cleft upon transition from the closed to the open state, providing the spatial capacity to accommodate the bulky peptidoglycan stem pentapeptide. Supporting this induced-fit model, molecular dynamics simulations indicate reduced backbone flexibility (lower RMSF values) of catalytic and substrate-interacting residues in the substrate-bound state, suggesting the stabilisation of a catalytically competent conformation.^20^

This dynamic gating behaviour aligns with an emerging structural paradigm within the M15 endopeptidase family, in which loop-mediated transitions between closed and open conformations regulate substrate accessibility. For example, in the homologous bacteriophage T5 endolysin (EndoT5), access to the catalytic cleft is governed by a regulatory calcium-binding loop that reduces intramolecular mobility and stabilises the enzyme in an ‘open’ conformation.

Together, these observations suggest that the Thr42–Ser75 loop acts as a dynamic gating element that regulates substrate access through a reversible closed-to-open conformational transition. However, because the simulations were performed using a minimal substrate model and the D156A mutant, these findings should be interpreted as indicative rather than definitive.^20,21^

### A divergent C-terminal helical element acts as a scaffold for substrate positioning

Structural and sequence comparison with representative M15 homologues showed that the C-terminal α-helix of LysPH is not conserved, suggesting that this element is a phage-specific adaptation. In contrast to some Gram-negative endolysin amphipathic helices that have been characterised, this helix does not exhibit features consistent with outer membrane disruption.^22–24^

Molecular dynamics simulations indicate that although the helix itself is not directly involved in contact with the glycan backbone, the adjacent loop region extends towards the active site cleft and makes transient contact with the peptidoglycan. These interactions guide the stem pentapeptide into a conformation that positions the scissile L-Ala–D-Glu bond proximal to the Zn^2+^ catalytic centre.

Substitution of Phe154 within this region increased local flexibility, indicating destabilisation of the C-terminal segment and supporting its structural role. Collectively, these results imply that the C-terminal helix is acting as a scaffold in the positioning of the adjacent loop region in such a way that the peptidoglycan is in the optimal position for the cross-linking peptide to access the catalytic site. ^25,26^

### Mechanistic model of catalysis and substrate positioning by LysPH

We propose a mechanistic model for LysPH activity based on structural, mutational, and computational analyses. In the absence of a substrate, LysPH adopts a closed conformation, in which the Thr42–Ser75 loop partially covers the catalytic cleft, restricting access to the active site, while the C-terminal region remains relatively flexible. Upon substrate engagement, the enzyme undergoes a substrate-induced transition to an open conformation; the flap region opens by hinge motion to accommodate the stem peptide of peptidoglycan within the active site. The C-terminal loop is seen to be in a transient interaction with the glycan backbone, positioning it so that the L-Ala-D-Glu bond is in proximity to the Zn^2+^ catalytic core.^27,28^

In the catalytically competent state, the Zn^2+^ ion and its coordinating residues polarise the scissile amide bond, promoting the activation of a bound water molecule for nucleophilic attack. Asp156 is proposed to stabilise the catalytic water network and/or facilitate substrate positioning, rather than acting as an essential catalytic base. In keeping with this model, mutations that disrupt Zn^2+^ coordination abolish enzymatic activity, while mutations that disrupt substrate-binding and positioning residues severely impair enzymatic activity.

### Implications for engineering and therapeutic development

The mechanistic insights obtained here provide a framework for the rational engineering of LysPH and related M15 endopeptidases, as the strong dependence of LysPH activity on peptidoglycan binding and the alignment of the cross-linking peptide within the active-site cleft indicates that modifications to stabilise the open conformation of the Thr42–Ser75 flap may enhance catalytic efficiency. Similarly, targeted alterations in the C-terminal loop could be used to tune substrate specificity and modulate host range.^8,29^

Notably, LysPH activity against Gram-negative strains was dependent on outer membrane permeabilisation, consistent with the general requirement for exogenous agents to enable globular endolysins to access the periplasmic peptidoglycan. In this respect, LysPH represents a promising candidate that can be developed to exhibit improved antimicrobial activity, perhaps through fusion with permeabilising peptides or by combining it with outer membrane permeabilising agents.

## Experimental procedures

### Bacteriophage isolation, purification and enrichment

Water samples were collected from different locations of the river Ganga, including Roorkee (29°52′29.49″N, 77°53′23.74″E), Haridwar (29°56′42″N, 78°09′47″E), Rishikesh (30°06′30″N, 78°17′50″E), and Varanasi (25°19′08″N, 83°00′46″E). Samples were centrifuged at 6,500 × g for 10 min, filtered through a sterile 0.22-μm syringe filter to remove sediment and endogenous bacteria.

Bacteriophages were isolated as previously described, with minor modifications. Briefly, 25 mL of filtered sample was mixed (1:1) with double-strength Luria-Bertani (LB) medium and inoculated with early-exponential-phase *P. aeruginosa DSMZ 50071* for phage enrichment. The mixture was incubated at 37 °C for 12 h with shaking at 200 rpm, followed by an additional incubation of 5 h at 4 °C. ^30–32^

After incubation, the mixture was centrifuged at 6,500 × g for 15 min to remove bacterial cells, and the supernatant was filtered again through a sterile 0.22-μm syringe filter. Phage detection and isolation were done using the double-layer agar method. Briefly, 100 μL of filtrate and 100 μL of *P. aeruginosa DSMZ 50071* were mixed with 3 mL of 0.7% (w/v) top LB agar (semi-solid) and overlaid onto LB agar plates, which were incubated for 8 h at 37 °C.

An isolated plaque was picked, resuspended in 100 μL of exponentially growing *P. aeruginosa*, and incubated for 4 h at 37 °C with shaking at 200 rpm. It was then mixed with 3 mL of top LB agar (semi-solid), poured onto LB agar plates, and incubated at 37 °C overnight. This step was repeated five times to ensure a homogeneous phage population. This phage was then designated PA_Ganga_001.

### DNA extraction, sequencing, and bioinformatic analysis

To eliminate host nucleic acid contamination, high-titer PA_Ganga_001 lysates (101^2^–101^4^ PFU mL^−1^) were incubated with DNase I and RNase A (1 µg mL^−1^ each) for 30 min at 37 °C. Phage genomic DNA was extracted as previously described, with minor modifications, using phenol:chloroform:isoamyl alcohol (25:24:1) followed by ethanol precipitation.^33^ DNA concentration and purity were assessed by spectrophotometry using a NanoDrop Eight Microvolume Spectrophotometer (Thermo Scientific, USA) and by agarose gel electrophoresis.

Paired-end whole-genome sequencing (2 × 150 bp, Illumina platform) was performed by a commercial sequencing provider, and raw FASTQ files were processed for downstream analysis. Adapter trimming and quality filtering were conducted using Cutadapt and Trimmomatic. High-quality reads were assembled de novo using SPAdes v3.15.5. De novo assembly produced a single contig of 63,392 bp with an average coverage of 100.2×. Open reading frames were predicted and annotated with Pharokka v1.7.1 and refined by BLASTp searches against the NCBI non-redundant database. The complete genome sequence of PA_Ganga_001 has been deposited in GenBank under accession number PP597238.^34–38^

### Phage stability and infection kinetics assays

The phage suspensions (10^8^ PFU mL^−1^ in SM buffer, pH 7.5) were incubated at 10-90 °C in 10 °C intervals for 1 h. The remaining infectivity was measured using the double-layer agar overlay assay, and the % of viable phages was calculated from the initial phage titre before the temperature treatment. For pH stability, phage suspensions (10^8^ PFU mL^−1^) were incubated at 37 °C for 1 h at pH 2–10 with Teorell–Stenhagen buffer (50 mM potassium chloride, 10 mM monopotassium phosphate, 10 mM trisodium citrate, and 10 mM boric acid) adjusted to the desired pH with NaOH and HCl, neutralised immediately after incubation, and titrated using the double-layer method. The percentage of viable phages at each pH was calculated relative to the initial phage titre before pH treatment. All assays were performed in triplicate.^39–41^

Adsorption kinetics of PA_Ganga_001 were determined using mid-exponential phase MDR *P. aeruginosa DSMZ 50071* (OD_600_ = 0.4) infected at a MOI of 0.01 in LB medium. At 0–50 min post-infection (5 min intervals), aliquots were withdrawn and immediately treated with chloroform to inactivate bacterial cells. The remaining free phage particles were quantified by the double-layer agar assay. The percentage of free phage particles remaining at each time point was calculated as 100 × (free phage titre/initial titre). Experiments were conducted in triplicate.^42–44^

A one-step growth assay was carried out as described earlier^45^, with some modifications. The mid-exponential phase culture of *P. aeruginosa DSMZ 50071* (OD_600_ ≈ 0.4) was infected with the phage at an MOI of 0.01 and incubated for 10 min at 37 °C. Cultures were centrifuged (6,000 × g, 10 min); the bacterial pellet was washed with fresh LB broth to eliminate unadsorbed phages, resuspended in pre-warmed LB broth to minimise secondary adsorption, and then diluted to a 1:100 ratio with LB. The bacterial culture was then incubated at 37°C with shaking at 200 rpm. Every 15 minutes, 50 μL of the sample was collected for 120 minutes, and each sample was then treated with chloroform, and the resulting suspension was quantified using the double-layer agar assay.

The titers of phage (PFU/mL) were graphed against time to obtain growth curves. The latent period was defined as the time between the end of adsorption and the onset of an increase in extracellular phage titers. The burst size was estimated as the ratio of the average phage titer during the plateau phase (∼9.2 × 10^9^ PFU mL^−1^) to the number of infected cells at time zero (∼10^10^ CFU mL^−1^). Each experiment was performed in triplicate assays.^45–47^

### Transmission electron microscopy

Phage morphology was examined by negative-stain transmission electron microscopy (TEM). Purified PA_Ganga_001 particles (107–10^9^ PFU mL^−1^) were applied to Formvar-coated 300-mesh copper grids and allowed to adsorb for 2 min at room temperature. The liquid was removed by blotting, and the grids were washed twice with double-distilled water. Negative staining was done using 2% (w/v) uranyl acetate for 60 seconds. Grids were analysed using a Talos F200X G2 S-TWIN (Thermo Fisher Scientific) transmission electron microscope operated at 200 kV at the Institute Instrumentation Centre (IIC), IIT Roorkee.^48^

### Cloning, expression and purification of LysPH and its mutants

The LysPH gene was amplified from genomic DNA of the lytic *P. aeruginosa* bacteriophage PA_Ganga_001 using gene-specific primers containing NdeI and HindIII restriction sites at the 5’ and 3’ ends, respectively. Amplified product and pET-28a (+) plasmid were digested, purified and ligated using T4 DNA ligase. Recombinant pET-28a-LysPH plasmids were transformed into chemically competent *E. coli BL21* (λDE3) for protein expression.

For protein expression, cells in a culture of 1 L of LB medium supplemented with kanamycin (50 μg mL^−1^) were induced at an OD_600_ of 0.6 using 1 mM isopropyl β-D-1-thiogalactopyranoside (IPTG) for 5h at 37°C.

Induced culture was harvested by centrifugation and resuspended in lysis buffer (50 mM Tris– HCl, pH 7.5; 250 mM NaCl; 15% glycerol). Cell disruption was performed using a constant cell disruptor at 24 kpsi, and the supernatant containing the LysPH was clarified by centrifugation. The supernatant was applied to a pre-equilibrated Ni-NTA affinity column, and the bound His-tagged LysPH protein was eluted using an imidazole gradient (100–500 mM) in purification buffer. Eluted fractions were analysed by SDS–PAGE. Protein samples were concentrated using a 10 kDa molecular weight cut-off centrifugal filter (Amicon, Millipore) and dialysed against storage buffer (50 mM HEPES, pH 7.5; 250 mM NaCl).^33^

Point mutations (H77A, H159A, D156A, D84A, R41A and S55A) were introduced into the LysPH expression construct by site-directed mutagenesis. Mutagenic primers carrying the desired nucleotide substitutions were designed according to standard overlap-extension principles. The parental LysPH-pET28a(+) plasmid was used as the template for whole-plasmid PCR amplification using a high-fidelity DNA polymerase.^49^

Following amplification, the PCR products were treated with DpnI to selectively digest the methylated parental plasmid DNA. The resulting nicked, mutated plasmids were directly transformed into chemically competent *Escherichia coli BL21 (λDE3)* cells. Transformants were selected on LB agar plates supplemented with kanamycin (50 μg mL^−1^). Mutations were confirmed by Sanger sequencing.^49^

All mutant proteins were expressed and purified using the same protocol described for wild-type LysPH.

Analytical size-exclusion chromatography (SEC) was performed to assess the folding, oligomeric state, and hydrodynamic properties of wild-type LysPH and alanine-substitution mutants (R41A, S55A, H77A, D84A, and H159A) using a HiLoad 16/600 Superdex 75 pg column (Cytiva) equilibrated in 50 mM HEPES (pH 7.5) and 250 mM NaCl. Protein samples were loaded under identical chromatographic conditions and eluted at a constant flow rate, with absorbance monitored at 280 nm. All constructs were analysed sequentially on the same column to ensure comparability. Minor differences in peak intensities (mAU) reflect variations in protein concentration at the time of injection rather than differences in oligomeric state, as all constructs eluted at comparable volumes. Chromatograms were processed in R following baseline correction and Savitzky–Golay smoothing, and peak elution volumes (Ve) were determined within the separation window. Elution behaviour was further evaluated using the partition coefficient (Kav) and peak shape parameters, confirming monodisperse profiles for all constructs.

### Turbidity reduction assay

Concentration-dependent activity measurements were performed using nine distinct concentrations of wild-type LysPH ranging from 0.01 to 50 µM. Turbidity reduction was monitored by recording the OD600 for 30 minutes. Data from three independent biological replicates (assayed in technical triplicate) were averaged and interpolated onto a common time grid (0.2 min steps). The endpoint % lysis was calculated at 30 minutes. Initial lysis rates were derived via linear regression of the OD600 time-course over the first 20 minutes. To calculate the half-maximal effective concentration (EC50) and Hill coefficients, both the endpoint % lysis and initial rate data were fitted to a four-parameter logistic (4PL) regression model using R.

Lytic activity of wild-type LysPH and six alanine-substitution mutants was assessed using a turbidity reduction assay as previously described. Five bacterial strains were tested: *A. baumannii (clinical isolate); K. pneumoniae 3796; K. pneumoniae 5362; K. pneumoniae (clinical isolate); P. aeruginosa DSM 50071*.

Bacteria were grown to mid-exponential phase, harvested by centrifugation, washed, and resuspended in reaction buffer (50 mM Tris-HCl, pH 8.0) to an OD_600_ of approximately 1.0. Gram-negative bacteria were pre-treated with 0.5 mM EDTA to permeabilise the outer membrane, as commonly required for globular endolysins to access periplasmic peptidoglycan. Cells were then washed three times with reaction buffer to remove residual EDTA before setting up the reactions.^50–52^

Reactions were performed in flat-bottom 96-well microplates in a final volume of 200 µL, comprising 100 µL of bacterial suspension and 100 µL of purified protein (2 µM stock, yielding a final concentration of 1 µM) or an equivalent volume of buffer control. OD_600_ was recorded every 5 seconds for 60 minutes using a microplate reader. For graphical representation, OD_600_ values were plotted at 10-minute intervals.

All experiments were performed in three independent biological replicates, each measured in three technical replicates (n = 3 biological replicates, with three technical replicates per biological replicate). Technical replicates were averaged prior to analysis, yielding n = 3 independent values per condition.^49,53^

Lytic activity was quantified as the area under the OD_600_ curve (AUC) over 60 minutes. For AUC calculations and graphical representation, data were down-sampled to 10-minute intervals to ensure uniform temporal spacing (Δt = 10 min) for trapezoidal integration. % OD reduction was calculated as:

% OD reduction = 100 × (1 − AUC_sample / AUC_buffer), where AUC_buffer is the mean AUC of wells receiving buffer instead of protein for the corresponding bacterial strain within the same experimental run.

For kinetic plots, OD values were normalised to the initial OD for each biological replicate (OD_t/OD_t=0), yielding a starting value of 1.0 for all conditions. For heatmap analysis, relative activity was calculated by dividing each mutant’s mean % OD reduction by the corresponding wild-type mean for the same strain (WT = 1.0).

Normality of data distributions was assessed using the Shapiro–Wilk test. For normally distributed data, statistical comparisons between each mutant and wild-type were performed using Welch’s t-test (unequal variance), whereas non-normal data were analysed using the Mann–Whitney U test. Multiple comparisons were corrected using the Holm method, applied separately for each bacterial strain. All analyses were performed using Python (scipy.stats, numpy, pandas) and independently validated in R (multcomp, DescTools).

### Crystallisation of LysPH and LysPH_D156A – pentapeptide bound complex

Purified LysPH and LysPH_D156A proteins were concentrated to 20 mg mL^−1^ and subjected to crystallisation screening using the sitting-drop vapour-diffusion method at 20 °C. Initial crystals of LysPH were obtained using the Hampton Research Crystal Screen 1. Optimisation yielded the best diffracting crystals in a reservoir solution containing 20% w/v Polyethylene glycol 3,350, 0.2 M Potassium sodium tartrate tetrahydrate, pH 7.4, with a protein-to-reservoir ratio of 1:1 (1 μL protein + 1 μL reservoir). Prior to data collection, crystals were briefly soaked in the reservoir solution containing 15% (v/v) ethylene glycol and subsequently flash-cooled in a nitrogen stream at 100 K.

To capture the enzyme–substrate complex, the D156A mutant was selected based on its proximity to, but not direct involvement in, Zn^2+^ coordination. Substitution of Asp156 was therefore expected to reduce catalytic turnover while preserving the integrity of the metal-binding site, thereby facilitating substrate trapping.

For LysPH_D156A, optimal crystals were obtained in a reservoir solution comprising 0.1 M sodium acetate trihydrate (pH 4.6), 0.2 M ammonium sulphate and 30% (w/v) Polyethylene glycol 2000, using a 1:1 protein-to-reservoir ratio at 20 °C. For ligand-bound complex formation, crystals were soaked for 15 min in reservoir solution supplemented with the Ala– D-γ-Glu–Lys–D-Ala–D-Ala peptide, which was obtained from Sigma-Aldrich (product code A0910, CAS 2614-55-3) at a concentration of 1 mM at 20 °C. Prior to data collection, crystals were briefly soaked in the reservoir solution containing 15% (v/v) ethylene glycol and subsequently flash-cooled in a nitrogen stream at 100 K.^54^

### X-ray data collection and structure determination

X-ray diffraction data were collected at the home source facility of the Macromolecular Crystallography Unit, IIC, IIT Roorkee, using a Rigaku MicroMax-007 HF microfocus rotating-anode X-ray generator equipped with a Rigaku HyPix 6000C detector. Diffraction images were indexed and integrated using CrysAlisPro (Agilent Technologies UK Ltd)^55^. Data scaling and merging were performed with AIMLESS in the CCP4i2 suite (v. 8.0.019)^56^. The structure was solved by molecular replacement using MOLREP in CCP4i2, with the coordinates of LysECD7 PDB 8Q2G used as the search model.^14^ Iterative cycles of manual model building and correction were performed in COOT, followed by restrained refinement with REFMAC in CCP4i2. For the pentapeptide-bound complex, difference Fourier maps (Fo– Fc) revealed clear electron density above 3σ, enabling modelling of the Ala–D-γ-Glu–Lys–D-Ala–D-Ala ligand into the active site. Omit maps were generated using PHENIX (v.1.18.2-3874)^57^ to validate ligand placement. Structural figures and interaction diagrams were prepared using UCSF ChimeraX (v. 1.8)^58^ and PyMOL (v. 2.5.5)^59,60^. Crystallographic data collection and refinement statistics for the LysPH and LysPH_D156A structures were generated using phenix.table_one and are summarised in Table 1.

**Table 1.**
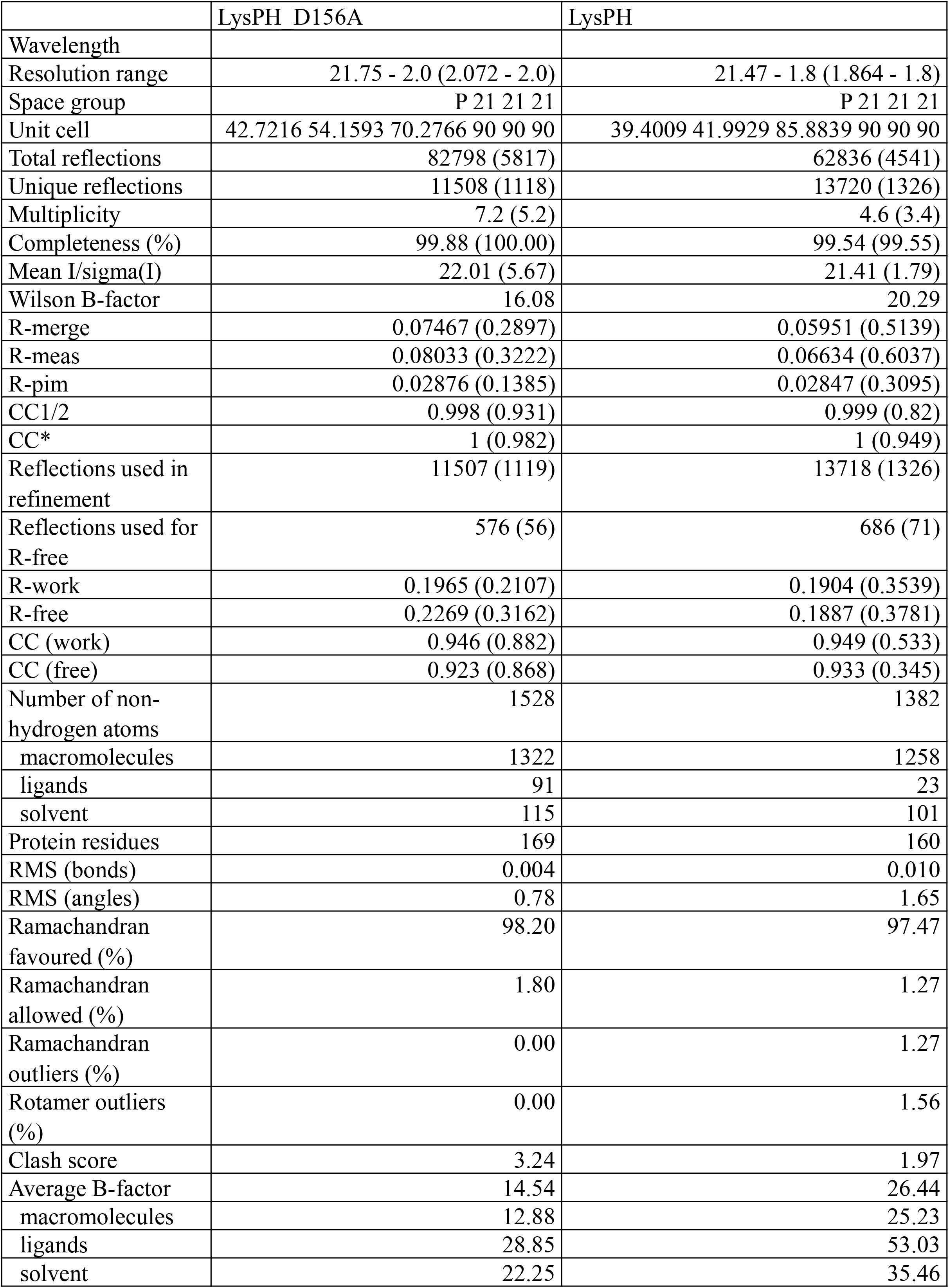
Data collection and refinement statistics.

|  | LysPH_D156A | LysPH |
| --- | --- | --- |
| Wavelength |  |  |
| Resolution range | 21.75 - 2.0 (2.072 - 2.0) | 21.47 - 1.8 (1.864 - 1.8) |
| Space group | P 21 21 21 | P 21 21 21 |
| Unit cell | 42.7216 54.1593 70.2766 90 90 90 | 39.4009 41.9929 85.8839 90 90 90 |
| Total reflections | 82798 (5817) | 62836 (4541) |
| Unique reflections | 11508 (1118) | 13720 (1326) |
| Multiplicity | 7.2 (5.2) | 4.6 (3.4) |
| Completeness (%) | 99.88 (100.00) | 99.54 (99.55) |
| Mean I/sigma(I) | 22.01 (5.67) | 21.41 (1.79) |
| Wilson B-factor | 16.08 | 20.29 |
| R-merge | 0.07467 (0.2897) | 0.05951 (0.5139) |
| R-meas | 0.08033 (0.3222) | 0.06634 (0.6037) |
| R-pim | 0.02876 (0.1385) | 0.02847 (0.3095) |
| CC1/2 | 0.998 (0.931) | 0.999 (0.82) |
| CC* | 1 (0.982) | 1 (0.949) |
| Reflections used in refinement | 11507 (1119) | 13718 (1326) |
| Reflections used for R-free | 576 (56) | 686 (71) |
| R-work | 0.1965 (0.2107) | 0.1904 (0.3539) |
| R-free | 0.2269 (0.3162) | 0.1887 (0.3781) |
| CC (work) | 0.946 (0.882) | 0.949 (0.533) |
| CC (free) | 0.923 (0.868) | 0.933 (0.345) |
| Number of non-hydrogen atoms | 1528 | 1382 |
| macromolecules | 1322 | 1258 |
| ligands | 91 | 23 |
| solvent | 115 | 101 |
| Protein residues | 169 | 160 |
| RMS (bonds) | 0.004 | 0.010 |
| RMS (angles) | 0.78 | 1.65 |
| Ramachandran favoured (%) | 98.20 | 97.47 |
| Ramachandran allowed (%) | 1.80 | 1.27 |
| Ramachandran outliers (%) | 0.00 | 1.27 |
| Rotamer outliers (%) | 0.00 | 1.56 |
| Clash score | 3.24 | 1.97 |
| Average B-factor | 14.54 | 26.44 |
| macromolecules | 12.88 | 25.23 |
| ligands | 28.85 | 53.03 |
| solvent | 22.25 | 35.46 |

### Molecular dynamics simulations

Molecular dynamics (MD) simulations were carried out for LysPH WT and the LysPH_D156A–pentapeptide complex using the Schrödinger Suite 2025-1^61^. Protein structures were built using the Protein Preparation Wizard in Maestro, where bond orders were assigned, side chains were added where necessary, hydrogen atoms were added, and protonation states were adjusted at pH 7.4. The OPLS4 force field was used for all simulations. Each system was solvated in an orthorhombic simulation box using the TIP3P water model, with a 1.0 nm buffer distance from the protein surface. Counter-ions were added to neutralise the system, and 0.15 M NaCl was added to simulate physiological ionic strength. System setup was carried out using the Desmond System Builder. Prior to production runs, the systems were subjected to a standard relaxation protocol comprising energy minimisation followed by stepwise equilibration under NVT and NPT ensembles to stabilise temperature and pressure. Simulations were run under NPT conditions for 200 ns at 300 K using the Nosé-Hoover thermostat and at 1 bar using the Martyna-Tobias-Klein barostat (relaxation time τ = 2.0 ps). A time step of 2 fs was used, and a 9.0 Å cutoff was used for short-range non-bonded interactions. Trajectory analyses, including root-mean-square deviation (RMSD), root-mean-square fluctuation (RMSF), and protein-ligand interaction analysis, were performed using the Simulation Interactions Diagram module in Desmond. All the simulations were run in triplicate.

## Acknowledgement

The authors gratefully acknowledge the financial support provided by the R&D department of THDC India Limited, Rishikesh, which made this research possible. We sincerely thank the R&D department and the organisation for its generous funding and continued encouragement towards scientific research and development. The X-Ray Crystallography facility was provided by MCU, IIC, IIT Roorkee. Ashok Soota Molecular Medicine Facility at the Department of Biosciences and Bioengineering, IIT Roorkee, was utilised for biochemical experimental work. The computational facility was provided by a DBT-funded project “Translational and Structural Bioinformatics” – BIC at the Department of Biotechnology, IIT, Roorkee (BT/PR40141/BTIS/137/16/2021) to P.K.

