## Supplementary Data for "Structural basis of substrate recognition and conformational gating in the bacteriophage M15 metalloendopeptidase LysPH"

### Isolation and characterisation of PA_Ganga_001

A lytic bacteriophage, PA_Ganga_001, was isolated from the Ganga River water samples using multidrug-resistant Pseudomonas aeruginosa DSMZ 50071 as the host bacterium. PA_Ganga_001 exhibited a podovirus-like morphology, with an isometric head and a short, non-contractile tail, characteristic of the class Caudoviricetes, as observed by transmission electron microscopy (Fig. S1).

PA_Ganga_001 is a double-stranded DNA phage with a genome size of 63,392 bp and a GC content of 55.95 %. Genomic annotation identified 115 coding sequences (CDSs), of which 85% encoded hypothetical proteins; the remaining 15% encoded structure and assembly proteins, including four connector proteins, eight head and packaging proteins, and nine tail assembly proteins. Additionally, three genes encoding proteins involved in lysis were identified, along with five genes involved in nucleotide metabolism. No genes related to lysogeny (integrases and excisionases) or antimicrobial resistance were found, further suggesting that PA_Ganga_001 is a lytic phage.

PA_Ganga_001 retained >60% infectivity across pH 6–10. At pH 7, after 1 h of incubation, ~90 % of the phage particles were recovered, whereas infectivity was lost below pH 5. Thermal stability assays showed that ~97 % of the phage particles remained stable up to 37 °C, but became unstable at temperatures ≥50 °C and could not recover at ≥70 °C. (Fig. S2 A, B)

In the adsorption assay, PA_Ganga_001 showed efficient adsorption to P. aeruginosa DSMZ 50071, with ~40% of the phage adsorbed within 20 min post-infection, and by 40 min, ~90% were adsorbed, approaching complete adsorption by 45 min. In the one-step growth curve assay, PA_Ganga_001 had a latent period of 45 min and a burst size of 92 PFU per infected cell, comparable to other lytic *P. aeruginosa* phages. (Fig. S2 C, D)

### Relative lytic activity of LysPH mutants studied as a heat map

To visualise mutant effects across all strains simultaneously, we generated a heatmap of relative activity normalised to wild-type (WT = 1.0) with hierarchical clustering (Fig. S3C). Clustering of mutant columns revealed a clear separation into three functional groups: (i) catalytic mutants D84A and H77A, together with the binding-site mutant R41A, formed a tightly clustered group with consistently severe loss of function across all strains (relative activity < 0.20 in 4/5 strains); (ii) H159A showed intermediate impairment (relative activity 0.10–0.20); and (iii) D156A clustered separately, retaining the highest residual activity (relative activity 0.14–0.66 depending on strain). The S55A mutant clustered with the severely impaired group; however, interpretation of its functional phenotype is complicated by its altered biophysical behaviour, suggesting that Ser55 may help maintain the structural integrity of the active-site region. This indicates that proper folding and local structural stability are critical for LysPH function.

Clustering of bacterial strains (rows) revealed that A. baumannii clinical strains showed the strongest discrimination between WT and mutant activity, with all six mutants exhibiting significantly reduced activity (all $p < 0.01$). K. pneumoniae clinical showed the weakest discrimination, with only R41A reaching statistical significance after correction ($p < 0.01$), likely reflecting the lower intrinsic susceptibility of this strain to LysPH (WT activity 23.1%)

and the correspondingly reduced dynamic range for detecting mutant-specific effects. P. aeruginosa DSM 50071 showed intermediate antibacterial activity discrimination, with all six mutants significantly different from WT ($p < 0.05$).

**MD simulation analysis of LysPH and LysPH_D156A**

To study the structural stability and substrate interaction of LysPH endopeptidase, molecular dynamics (MD) simulations were performed for 200 ns on the apo wild-type enzyme and the substrate-bound LysPH_D156A mutant complexed with the peptidoglycan pentapeptide. Following an initial equilibration phase, both systems exhibited stable root-mean-square deviation (RMSD) relative to the apo enzyme (3.26 ± 0.54 Å) and the LysPH_D156A–pentapeptide complex (2.63 ± 0.32 Å), indicating overall structural stability throughout the simulation. The root-mean-square fluctuation (RMSF) of Cα atoms for catalytic residues and residues involved in substrate positioning decreases in the pentapeptide complex, indicating reduced backbone flexibility, with the mean RMSF of these residues decreasing from 1.20 Å to 0.67 Å. This reduction in RMSD and RMSF in the pentapeptide-bound mutant complex indicates increased structural stabilisation upon substrate binding, which may facilitate substrate positioning for catalysis (Fig. 5).

**Structural contribution of Ser55 to LysPH stability and function**

The only mutant exhibiting anomalous biophysical properties is the S55A mutant, which showed an abnormal SEC profile with significant structural perturbation. Unlike other mutants, which showed SEC profiles similar to those of the wild-type, with monodisperse species, the S55A mutant showed an abnormal shift in elution volume beyond the expected separation range ($K_{av} > 1.0$), with pronounced peak broadening and leading asymmetry. This abnormal elution is characteristic of non-specific interactions with the chromatographic matrix, which is usually due to exposed hydrophobic surfaces. In addition, increased peak width indicates conformational heterogeneity or the presence of multiple species in solution, whereas leading asymmetry indicates a subpopulation with an expanded hydrodynamic radius.

These observations indicate that substituting Ser55 affects the structural integrity of LysPH. Ser55 is located in the substrate-binding cleft of LysPH. This residue may be part of a local hydrogen bonding network that stabilises the active site architecture. Substitution of Ser55 with alanine is likely to alter this local environment, leading to partial unfolding or increased conformational flexibility.

A)

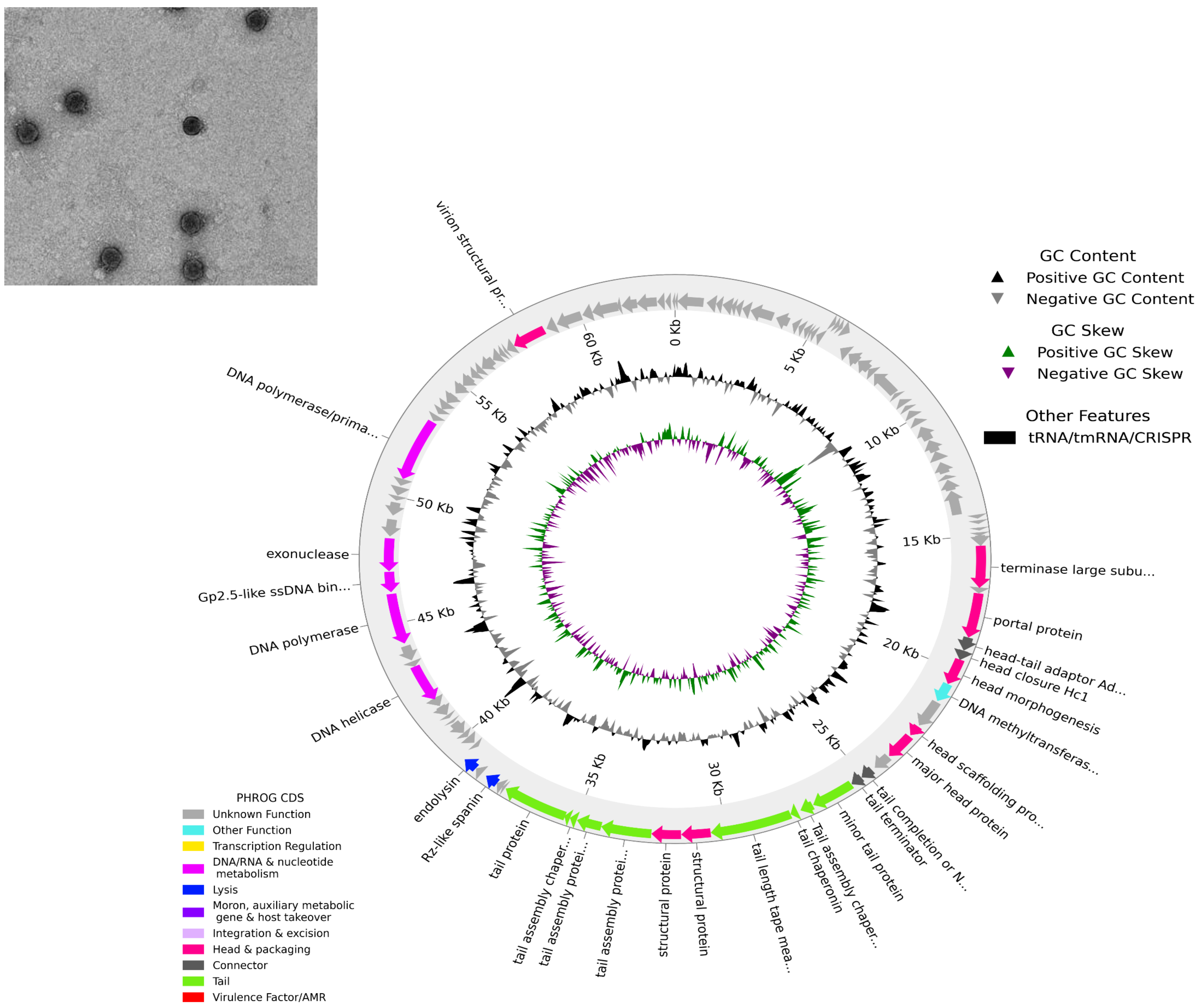


B)

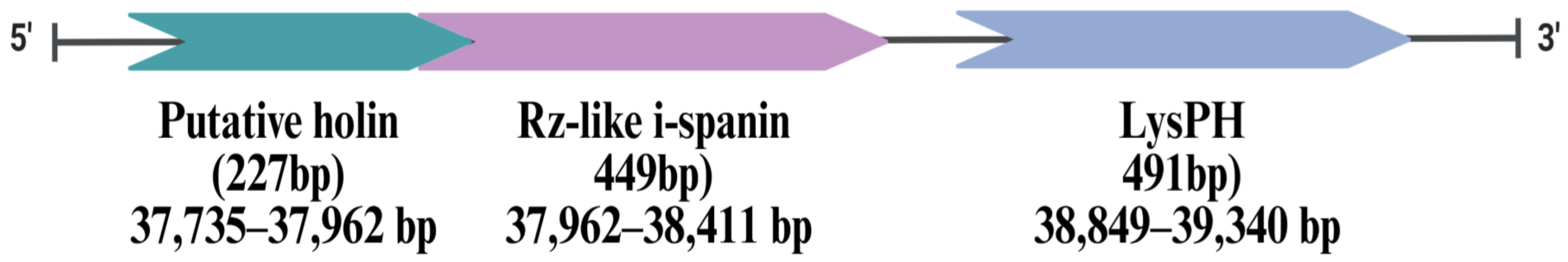


Figure S1: A) Transmission electron micrograph of negatively stained PG001 showing an isometric capsid with podovirus-like morphology. Scale bar, 100 nm. Circular genome map of PG001 (63 kbp; 55.9% GC; 115 predicted coding sequences). Genes are colour-coded by functional category (replication, structural/assembly, lysis, and hypothetical proteins). B) ) Lysis cassette of bacteriophage PG001.

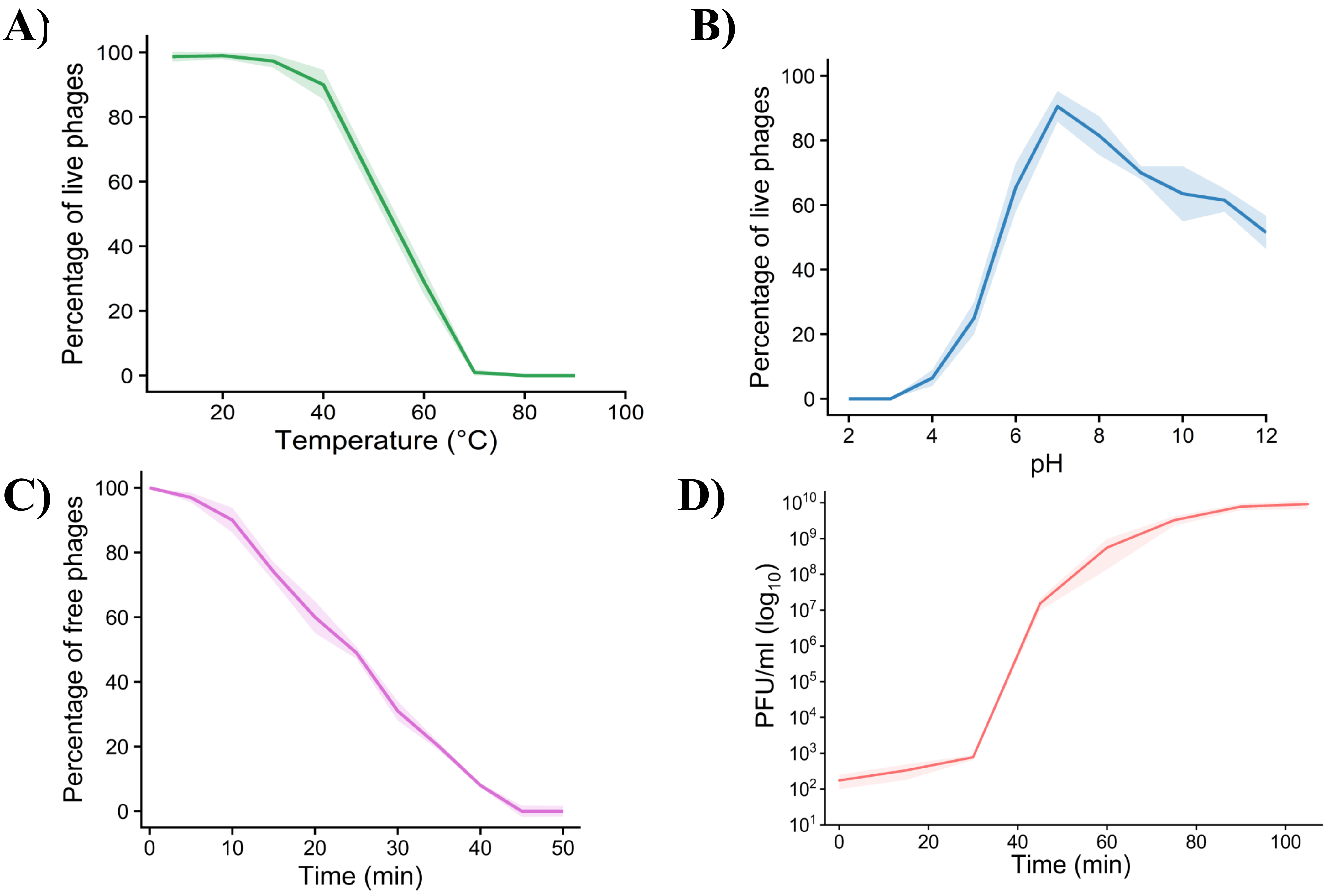


Figure S2: Biological characterisation of PG001 (A) Thermal stability of PG001 following 1 h incubation at the indicated temperatures. (B) pH stability of PG001 following 1 h incubation in buffers adjusted to the indicated pH values. (C) Adsorption kinetics of PG001 on multidrug-resistant *Pseudomonas aeruginosa* at MOI 0.01. (D) One-step growth curve of PG001 at MOI 0.01, showing latent and rise phases.

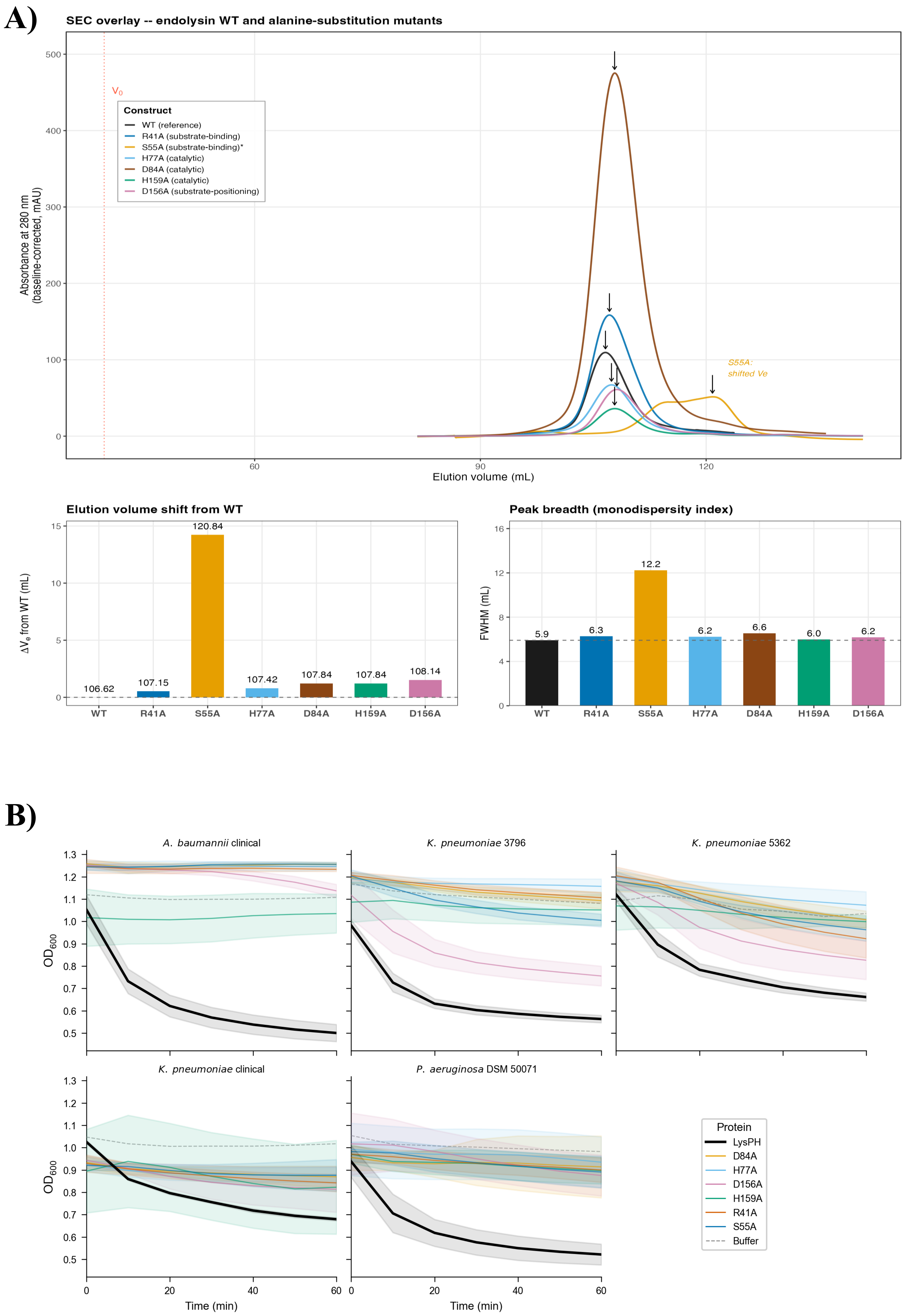


Figure S3: Biochemical characterisation of LysPH and its mutants (A) Time-course $OD_{600}$ reduction assay showing that wild-type LysPH (10 µM) exhibits rapid lysis across five Gram-negative strains, whereas catalytic (D84A, H77A, H159A), positioning (D156A), and binding-site (R41A, S55A) mutants display markedly reduced activity (mean ± SD, n = 3). Biophysical validation of LysPH and confirmation of structural integrity for catalytic mutants. SEC analysis of endolysin WT and five alanine-substitution mutants. Data: baseline correction (median 60-80 mL), Savitzky-Golay smoothing (window = 51, poly = 3), peak detection within 90-130 mL. Column: HiLoad 16/60

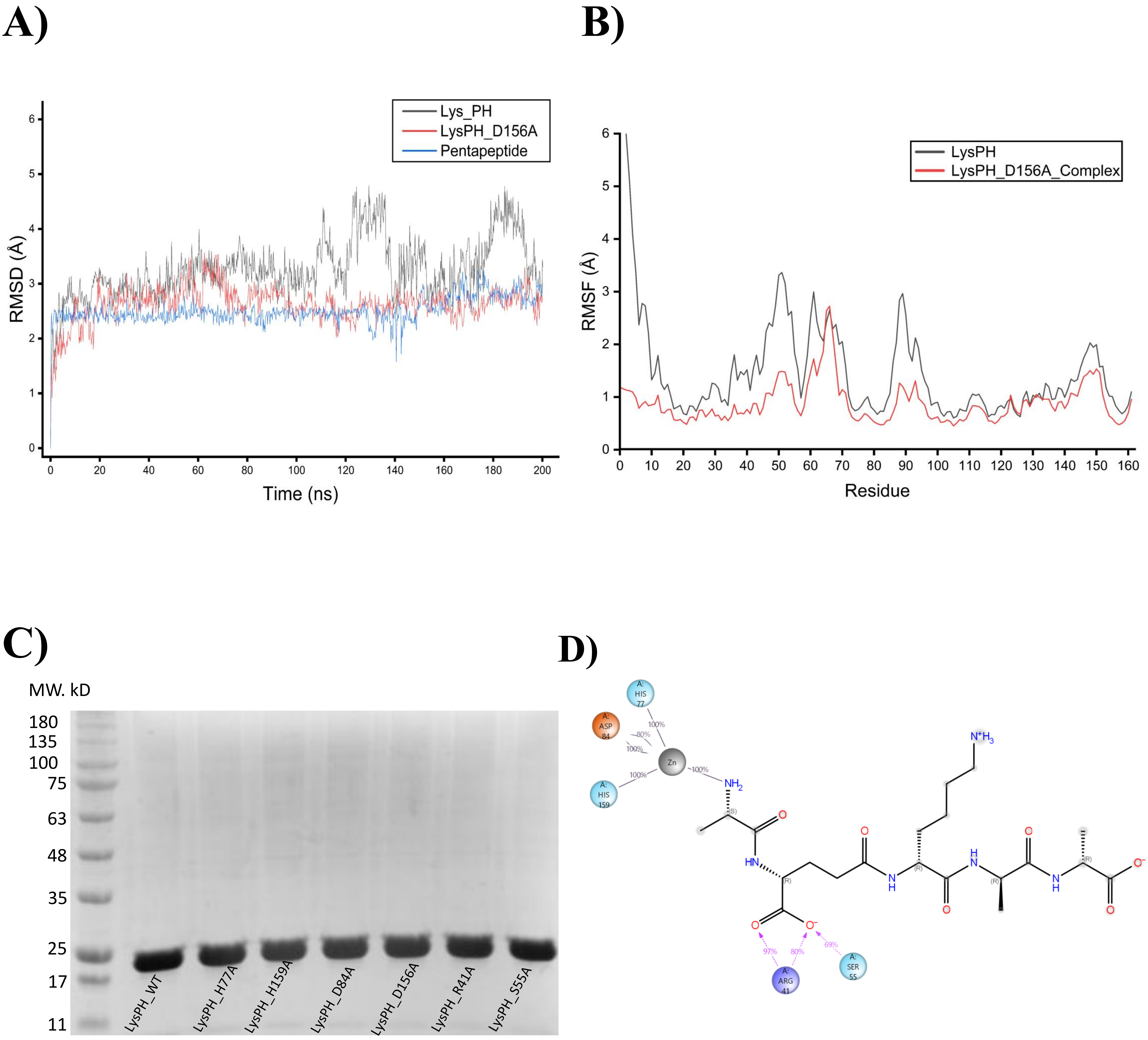


Figure S4. Molecular dynamics confirms substrate-induced active-site stabilisation and dynamic structural positioning. (A) Root mean square deviation (RMSD) of the LysPH–pentapeptide complex during the MD simulation, indicating structural stability. (B) Root mean square fluctuation (RMSF) of LysPH residues showing flexibility across the protein structure. (C) SDS–PAGE of purified LysPH and mutants (D) Ligand–protein interaction diagram generated from trajectory analysis showing contacts between the pentapeptide substrate and residues lining the catalytic groove of LysPH. Percentages indicate interaction occupancy during the simulation.

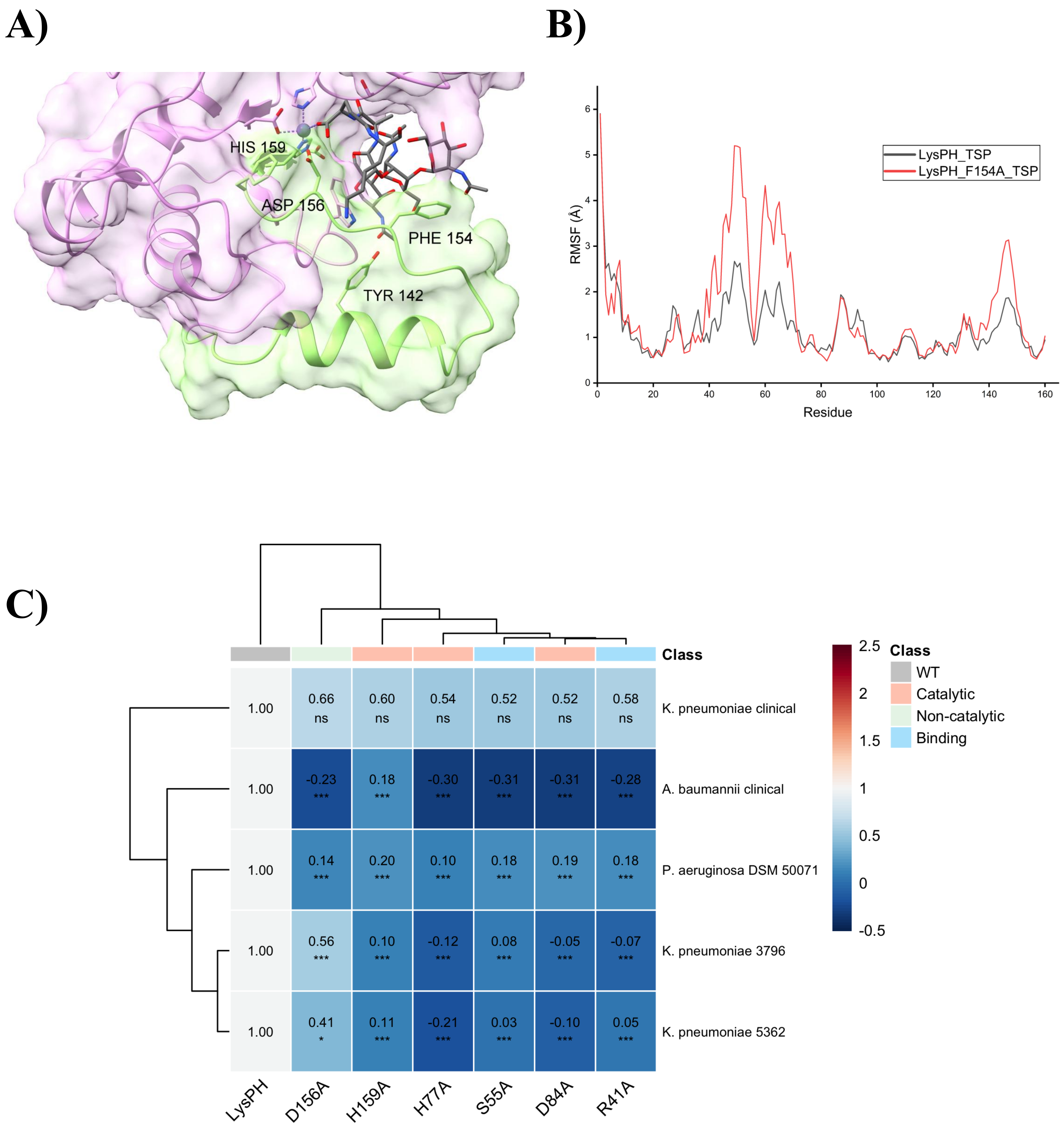


Figure S5. Molecular dynamics analysis of LysPH and LysPH_F154A with NAG–NAM–NAG–L-Ala–D-Glu–L-Lys. (A) C-terminal with substrate showing positioning of substrate within the catalytic site (B) Root mean square fluctuation (RMSF) of LysPH and LysPH_F154A residues showing flexibility across the protein structure. C) Relative lytic activity of LysPH mutants displayed as a heat map.

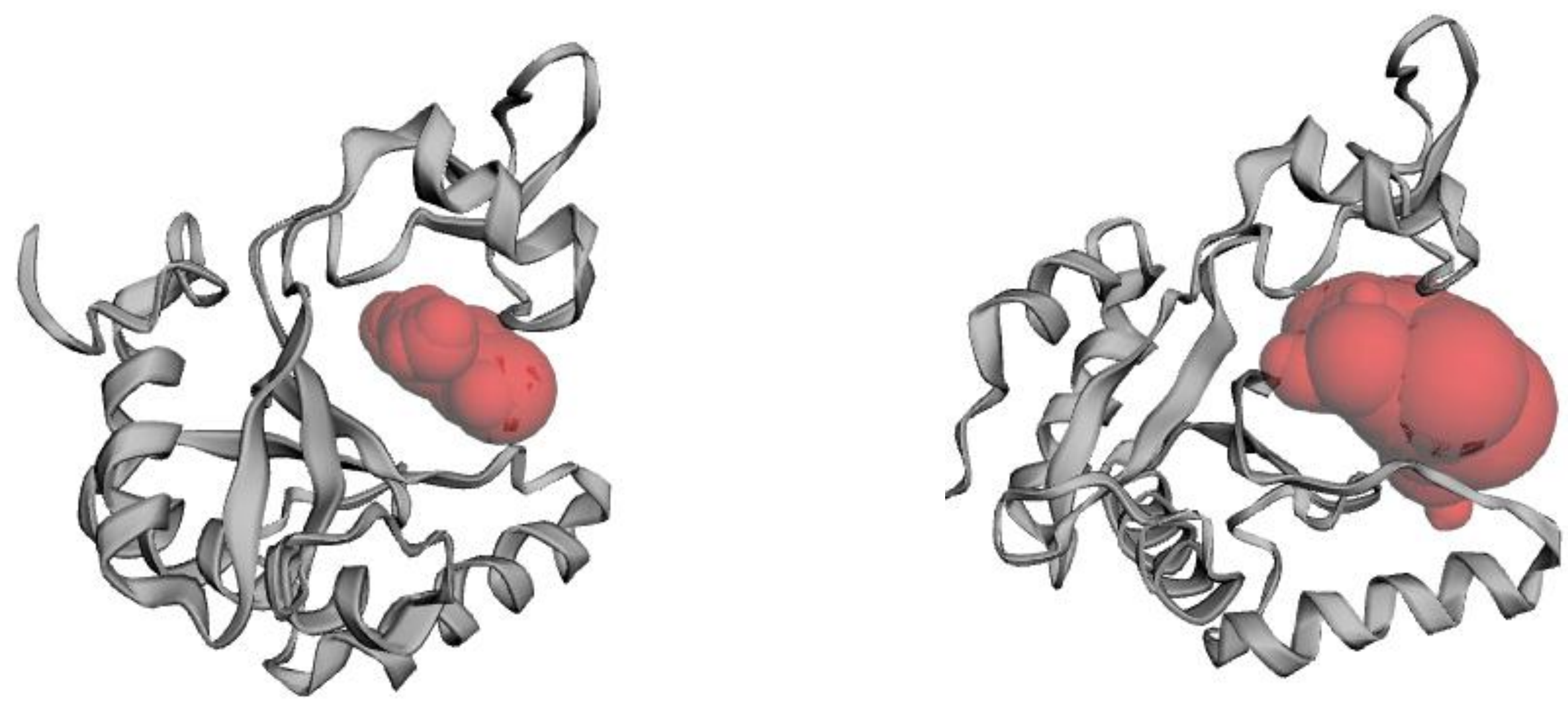

Figure S6. The solvent-accessible volume of the catalytic cleft of (A) LysPH and (B) LysPH_D156A measured using CastPfold

**Table S1.** Antibacterial efficacy of wild-type LysPH and active-site variants against Gram-negative pathogens.

| Protein | A. baumannii clinical | K. pneumoniae 3796 | K. pneumoniae 5362 | K. pneumoniae clinical | P. aeruginosa DSM 50071 |
|---|---|---|---|---|---|
| LysPH (WT) | 43.3 +/- 4.8 (1.00) | 41.7 +/- 1.8 (1.00) | 26.6 +/- 2.8 (1.00) | 23.1 +/- 0.4 (1.00) | 38.4 +/- 1.7 (1.00) |
| D84A | -13.3 +/- 0.7 (-0.31) ** | -2.2 +/- 1.8 (-0.05) *** | -2.5 +/- 2.7 (-0.10) ** | 12.1 +/- 2.8 (0.52) ns | 7.4 +/- 8.7 (0.19) * |
| H77A | -13.1 +/- 1.7 (-0.30) ** | -5.0 +/- 2.0 (-0.12) *** | -5.5 +/- 4.3 (-0.21) ** | 12.5 +/- 2.8 (0.54) ns | 3.7 +/- 4.1 (0.10) ** |
| H159A | 7.6 +/- 8.4 (0.18) ** | 4.1 +/- 4.4 (0.10) ** | 3.0 +/- 6.5 (0.11) * | 13.9 +/- 21.2 (0.60) ns | 7.8 +/- 5.1 (0.20) ** |
| D156A | -9.8 +/- 0.3 (-0.23) ** | 23.2 +/- 5.4 (0.56) * | 11.0 +/- 8.1 (0.41) ns | 15.3 +/- 7.0 (0.66) ns | 5.3 +/- 3.9 (0.14) ** |
| R41A | -12.1 +/- 1.6 (-0.28) ** | -3.1 +/- 2.0 (-0.07) *** | 1.2 +/- 6.5 (0.05) * | 13.5 +/- 0.9 (0.58) ** | 7.1 +/- 4.4 (0.18) * |
| S55A | -13.4 +/- 0.9 (-0.31) ** | 3.1 +/- 2.6 (0.08) *** | 0.8 +/- 1.9 (0.03) ** | 12.0 +/- 3.8 (0.52) ns | 6.8 +/- 4.6 (0.18) ** |

*Values shown as: % OD reduction +/- SD (relative activity) significance. Relative activity = mutant mean / WT mean for the same strain. * $p < 0.05$, ** $p < 0.01$, *** $p < 0.001$, ns = not significant.*

**Table S2.** Statistical analysis of mutational effects on LysPH antibacterial activity.

| Strain | Mutant | Delta (%) | p_adj | sig |
|---|---|---|---|---|
| A. baumannii clinical | D84A | 56.61938 | < 0.001 | *** |
| A. baumannii clinical | H77A | 56.33693 | < 0.001 | *** |
| A. baumannii clinical | H159A | 35.65817 | < 0.001 | *** |
| A. baumannii clinical | D156A | 53.04153 | < 0.001 | *** |
| A. baumannii clinical | R41A | 55.41745 | < 0.001 | *** |
| A. baumannii clinical | S55A | 56.70318 | < 0.001 | *** |
| K. pneumoniae 3796 | D84A | 43.93693 | < 0.001 | *** |
| K. pneumoniae 3796 | H77A | 46.7509 | < 0.001 | *** |
| K. pneumoniae 3796 | H159A | 37.62662 | < 0.001 | *** |
| K. pneumoniae 3796 | D156A | 18.51705 | < 0.001 | *** |
| K. pneumoniae 3796 | R41A | 44.86854 | < 0.001 | *** |
| K. pneumoniae 3796 | S55A | 38.60348 | < 0.001 | *** |
| K. pneumoniae 5362 | D84A | 29.07881 | < 0.001 | *** |
| K. pneumoniae 5362 | H77A | 32.06391 | < 0.001 | *** |
| K. pneumoniae 5362 | H159A | 23.55837 | < 0.001 | *** |
| K. pneumoniae 5362 | D156A | 15.53942 | 0.012 | * |
| K. pneumoniae 5362 | R41A | 25.32527 | < 0.001 | *** |
| K. pneumoniae 5362 | S55A | 25.78849 | < 0.001 | *** |
| K. pneumoniae clinical | D84A | 10.95734 | 0.471 | ns |
| K. pneumoniae clinical | H77A | 10.59724 | 0.502 | ns |
| K. pneumoniae clinical | H159A | 9.171717 | 0.632 | ns |
| K. pneumoniae clinical | D156A | 7.791975 | 0.759 | ns |
| K. pneumoniae clinical | R41A | 9.601721 | 0.592 | ns |
| K. pneumoniae clinical | S55A | 11.04076 | 0.464 | ns |
| P. aeruginosa DSM 50071 | D84A | 30.9643 | < 0.001 | *** |
| P. aeruginosa DSM 50071 | H77A | 34.65402 | < 0.001 | *** |
| P. aeruginosa DSM 50071 | H159A | 30.57887 | < 0.001 | *** |
| P. aeruginosa DSM 50071 | D156A | 33.04768 | < 0.001 | *** |
| P. aeruginosa DSM 50071 | R41A | 31.30881 | < 0.001 | *** |
| P. aeruginosa DSM 50071 | S55A | 31.54641 | < 0.001 | *** |

Differences in lytic activity (Δ, expressed as percent $OD_{600}$ reduction relative to wild-type) were calculated for each mutant across all tested strains. Statistical significance was assessed using Welch's *t*-test with Holm correction for multiple comparisons. Adjusted *p*-values ($p_{adj}$) and corresponding significance levels are reported as ns (not significant), * ($p < 0.05$), ** ($p < 0.01$), and *** ($p < 0.001$). Data indicate a consistent and significant loss of activity for most mutants across susceptible Gram-negative strains, whereas no statistically significant differences were observed for *K. pneumoniae* clinical, consistent with its low intrinsic susceptibility to LysPH.
